# Noninvasive reconstruction of complete mitochondrial genomes from aquatic environmental DNA using PCR-free long-read sequencing

**DOI:** 10.1101/2025.06.24.661187

**Authors:** Hinano Mizuno, Hidenori Tanaka, Yoshikazu Furuta, Nobuhiko Muramoto

**Author notes:** **Corresponding author** Hinano Mizuno.

## Abstract

1. Environmental DNA (eDNA) is a powerful, noninvasive data source for assessing aquatic biodiversity, but the assumption that it is present only in low concentrations and highly fragmented forms has limited its use. Accordingly, eDNA genomic analyses have focused on short mitochondrial regions, which often lack the resolution for accurate species identification or individual-level analysis. However, some intact mitochondrial genomes may persist in aquatic environments, and their recovery could enable noninvasive determination of complete mitochondrial sequences, providing higher-resolution genetic information. Nevertheless, most eDNA studies have overlooked long-read DNA analysis, and current protocols are not optimised to detect or utilise these longer fragments.
2. To address this, we developed a long-read sequencing workflow to reconstruct complete mitochondrial genomes from eDNA. First, we selected the optimal DNA extraction method and filter pore sizes using water from aquaria containing *Carassius auratus* or *Oryzias latipes*, aiming to maximise the yield of high-molecular-weight DNA for target organism sequences. We then applied two sequencing strategies: a PCR-free approach using native eDNA and a long-range PCR-based approach targeting the complete mitochondrial genome.
3. Using the determined DNA extraction method and a filter (1.2 µm pore size), we obtained high-molecular-weight DNA suitable for long-read sequencing. In the PCR-free approach, *C. auratus* samples from single-fish aquaria yielded hundreds of reads mapped to the mitochondrial genome, with some reads exceeding 16,000 bp. These allowed *de novo* assembly of full-length mitochondrial genomes in several samples. In the PCR-based approach, one sample yielded over 100,000 mitochondrial reads, of which >27% exceeded 16,000 bp, enabling assemblies. Comparison with assemblies from tissue-derived DNA confirmed accuracy, as sequences were nearly identical. Across samples, polymorphism analysis revealed two distinct haplotypes in *C. auratus*. These results demonstrate that full-length mitochondrial genomes can be reconstructed from eDNA, and that both PCR-free and PCR-based strategies are effective under appropriate sample conditions.
4. Our results show that long-read sequencing enables the recovery of near-complete mitochondrial genomes directly from eDNA without requiring artificial fragmentation and, in some cases, without the need for enrichment steps. This approach enhances eDNA-based biodiversity monitoring by improving species identification and enabling within-population level analyses.

## 1. Introduction

Biodiversity plays a crucial role in providing essential services, such as food, water, and medicinal resources and it supports human health by regulating ecosystems (Aerts et al., 2018; Li et al., 2024). The loss of biodiversity can compromise these functions and negatively impact human well-being (Zhang et al., 2022). Biodiversity refers to the variety of ecosystems, species, and genetic diversity that collectively underpin ecological stability. This includes genetic diversity, which is vital for supporting ecological resilience and adaptability (Fargeot et al., 2024; Stange et al., 2021; Roger et al., 2012; Frankham, 1996). As the importance of genetic diversity becomes increasingly recognised, interest in this topic has grown (Frankham, 2015; Manel & Holderegger, 2013; Mable, 2019). However, conventional genetic assessments often require invasive tissue sampling, posing ethical and logistical challenges, especially for endangered or remote species (Taberlet et al., 1999; Thomsen & Willerslev, 2015). Therefore, developing noninvasive yet reliable alternatives is critical for conservation and population management.

Environmental DNA (eDNA), originating from organisms and present in the environment, has emerged as a promising noninvasive source for genetic analysis (Deiner et al., 2017a). It enables access to genetic information without sampling individuals, facilitating biodiversity assessments across ecosystems (Beng & Corlett, 2020). Most eDNA studies target mitochondrial regions to identify species, but understanding intraspecific diversity is also vital for conservation and evolutionary biology. Recent studies have applied eDNA in such analyses (Sigsgaard et al., 2020), using species-specific primers for phylogeographic investigations (Tsuji et al., 2023), comparing microsatellite-based allele frequencies (Andres et al., 2021), and estimating genotype frequency via cycling probe techniques (Uchii et al., 2016). These approaches typically amplify short sequences (100–500 bp), limiting their scope (Andres et al., 2023).

Analysing longer sequences, like complete mitochondrial genomes, can provide deeper insights into intraspecific phylogenetics, evolutionary history, and population structure (Miya et al., 2003; Das et al., 2025). However, since eDNA is generally highly fragmented (Hassan et al., 2022), it is unlikely that full-length mitochondrial DNA has been directly recovered from environmental samples without fragmentation during extraction or library preparation. Nonetheless, some indirect studies have suggested that such intact mitochondrial DNA may persist in aquatic environments (Moushomi et al., 2019; Deiner et al., 2017b).

Long-read sequencing (LRS), which has advanced rapidly, could enable recovery of intact full-length mitochondrial DNA from eDNA. A key advantage is its ability to sequence entire mitochondrial genomes as continuous strands, assuming intact DNA is present (Warburton & Sebra, 2023). This can aid in haplotype phasing (Browning & Browning, 2011), structural variant detection (Chaisson et al., 2019), and, without PCR, capture individual-level haplotypes and rare variants such as heteroplasmy (Sethi et al., 2019; Slapnik et al., 2024).

However, the use of LRS for intraspecific genetic diversity analysis from eDNA has not yet been reported, and its feasibility and utility remain untested. While several studies have used LRS for eDNA-based species detection (Nousias et al., 2024; Egeter et al., 2022; Doorenspleet et al., 2025), none aimed to recover full mitochondrial genomes of target species. Thus, it is necessary to establish a method to recover the complete mitochondrial genomes of target organisms from eDNA and to evaluate its applicability to genetic diversity analysis.

Here, we developed a workflow to recover full-length mitochondrial genomes from eDNA using LRS (Fig. 1). We optimised preprocessing and library preparation before implementing a sequence and analysis pipeline. This noninvasive approach enables eDNA-based intraspecific diversity analysis, offering applications in conservation, population management, and biodiversity monitoring.

**Figure 1.**
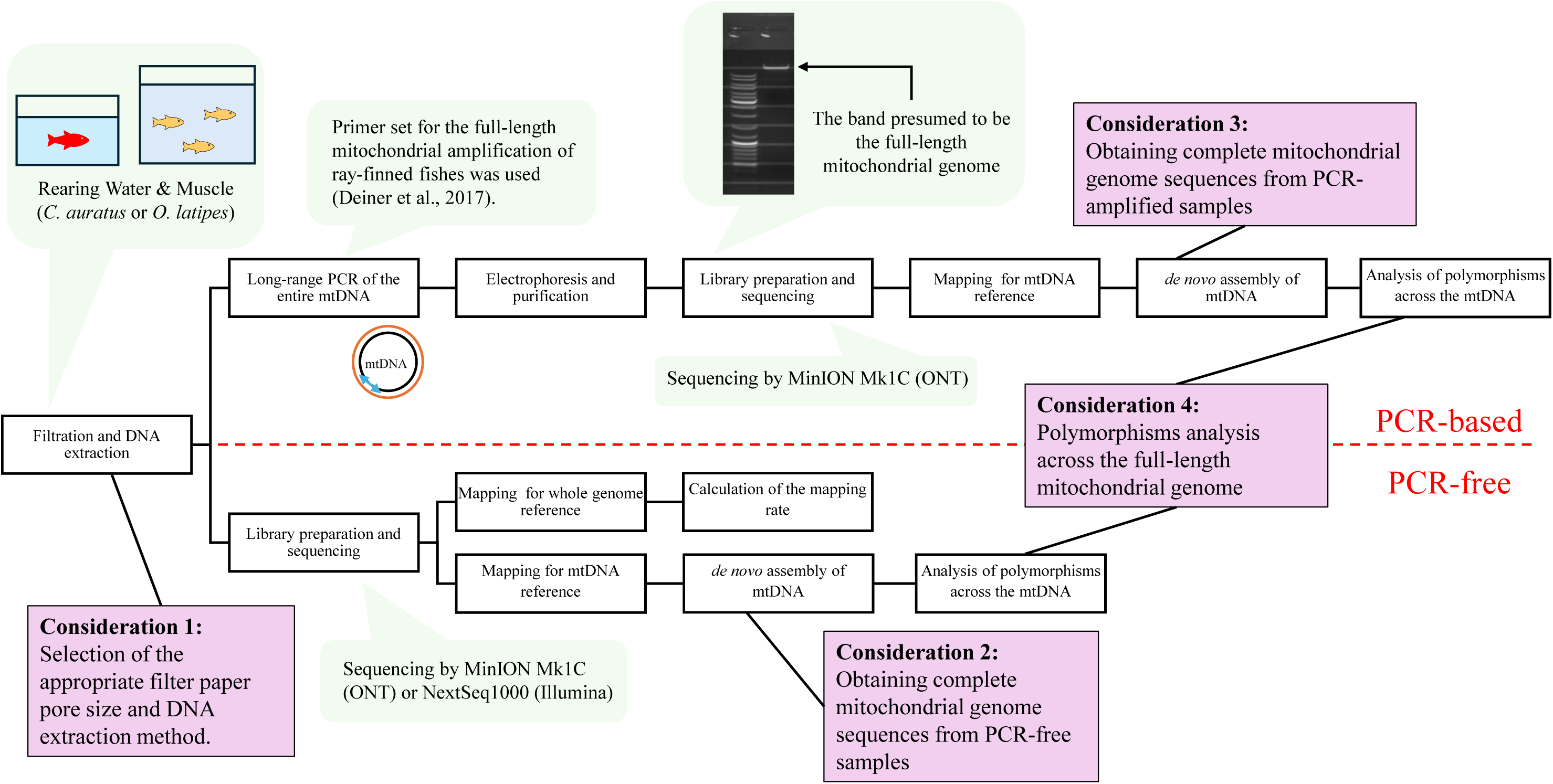
Experimental and analytical workflows of full-length mitochondrial amplification: PCR-based vs. PCR-free approaches A schematic diagram illustrating the two strategies we developed in this study for obtaining full-length mitochondrial genomes from eDNA: a PCR-based and a PCR-free workflow. Includes considerations for filtration, extraction, amplification, sequencing, and assembly steps.

## 2. Materials and Methods

### 2.1. Optimising the conditions for extracting high-molecular-weight DNA and recovering target species DNA

#### 2.1.1. Sample collection and DNA extraction

We used water from aquaria containing goldfish (*Carassius auratus*) or Japanese rice fish (*Oryzias latipes*) to optimise the conditions for recovering long DNA from target organisms, as these species were easy to obtain and maintain in laboratory aquaria. *C. auratus* (20–30 cm) were kept in a 100 L tank containing 14–22 fish, and *O. latipes* (2– 3 cm) in a 300 L tank containing 30–40 fish. From each tank, 250–1,000 mL of water was collected and immediately vacuum-filtered using 47 mm polycarbonate membrane filters with pore sizes of 0.2, 0.8, 1.2, 2.0, 3.0, 5.0, or 10.0 µm (Isopore, MilliporeSigma, Burlington, MA, USA).

DNA was extracted from the filters using the Blood & Cell Culture DNA Mini Kit (QIAGEN, Hilden, Germany) following the manufacturer’s instructions. The concentration of each extracted DNA solution was measured using the Qubit® 2.0 Fluorometer (Thermo Fisher Scientific, Waltham, MA, USA). The fragment lengths were examined using Genomic DNA ScreenTape, Genomic DNA reagents, and a 4150 TapeStation system (Agilent, Santa Clara, CA, USA). We also measured the bacterial DNA concentration using the Femto Bacterial DNA Quantification Kit (Zymo Research, Irvine, CA, USA) and the fungal DNA concentration using the Femto Fungal DNA Quantification Kit (Zymo Research), both on a QuantStudio® 3 Real-Time PCR System (Thermo Fisher Scientific).

In addition, the DNeasy Blood & Tissue Kit (QIAGEN), Extractor® WB Kit (FUJIFILM Wako Pure Chemical, Osaka, Japan), and MagAttract HMW DNA Kit (QIAGEN) were used to extract DNA from *C. auratus* aquarium water to compare fragment lengths and yields under different extraction procedures.

#### 2.1.2. Sequencing and bioinformatics

Libraries were prepared using the Ligation sequencing gDNA-Native Barcoding Kit 24 V14; SQK-NBD114.24 or the Ligation sequencing gDNA; SQK-LSK109 protocol (Oxford Nanopore Technologies, Oxford, UK). Sequencing was subsequently performed on a MinION Mk1C (Oxford Nanopore Technologies) equipped with FLO-MIN114, R10.4.1, or FLO-MIN106, R9.4.1 flow cells (Oxford Nanopore Technologies) following the manufacturer’s instructions. The pod5 and fast5 files were basecalled into fastq using Dorado v0.7.2 (Oxford Nanopore PLC, 2024) and guppy v1.1.alpha13 (Oxford Nanopore Technologies, 2024), respectively. Adapter trimming was conducted using Porechop v0.2.4 (Wick, 2024) and Chopper v0.8.0 (De Coster, 2024). Trimmed reads were mapped to the *C. auratus* (GCF_003368295.1) or *O. latipes* (GCF_002234675.1) reference genomes using Minimap2 v2.28 (Li, 2018) to generate BAM files. Mapping rates were calculated using SAMtools v1.21 (Danecek et al., 2021) and visualised with IGV (Robinson et al., 2011) and plot-bamstats (Torreso, 2024).

### 2.2. Retrieval of the complete mitochondrial sequence under different rearing conditions

#### 2.2.1. Sample collection and DNA extraction

Water was collected from aquaria in which *C. auratus* or *O. latipes* were reared under varying conditions (population size, tank volume, and duration) to obtain mitochondrial sequences (Table 1, Groups 1–6). From each tank, 250–1,000 mL was vacuum-filtered using 3.0 µm (Group 1) or 1.2 µm (Groups 2–6) membrane filters. DNA was extracted using the Blood & Cell Culture DNA Mini Kit and analysed for concentration and fragment length. Additionally, DNA was extracted from the muscle tissue of *C. auratus* or *O. latipes* individuals that died naturally (e.g. by jumping or illness) (Table 1, Group 7).

**Table 1.**
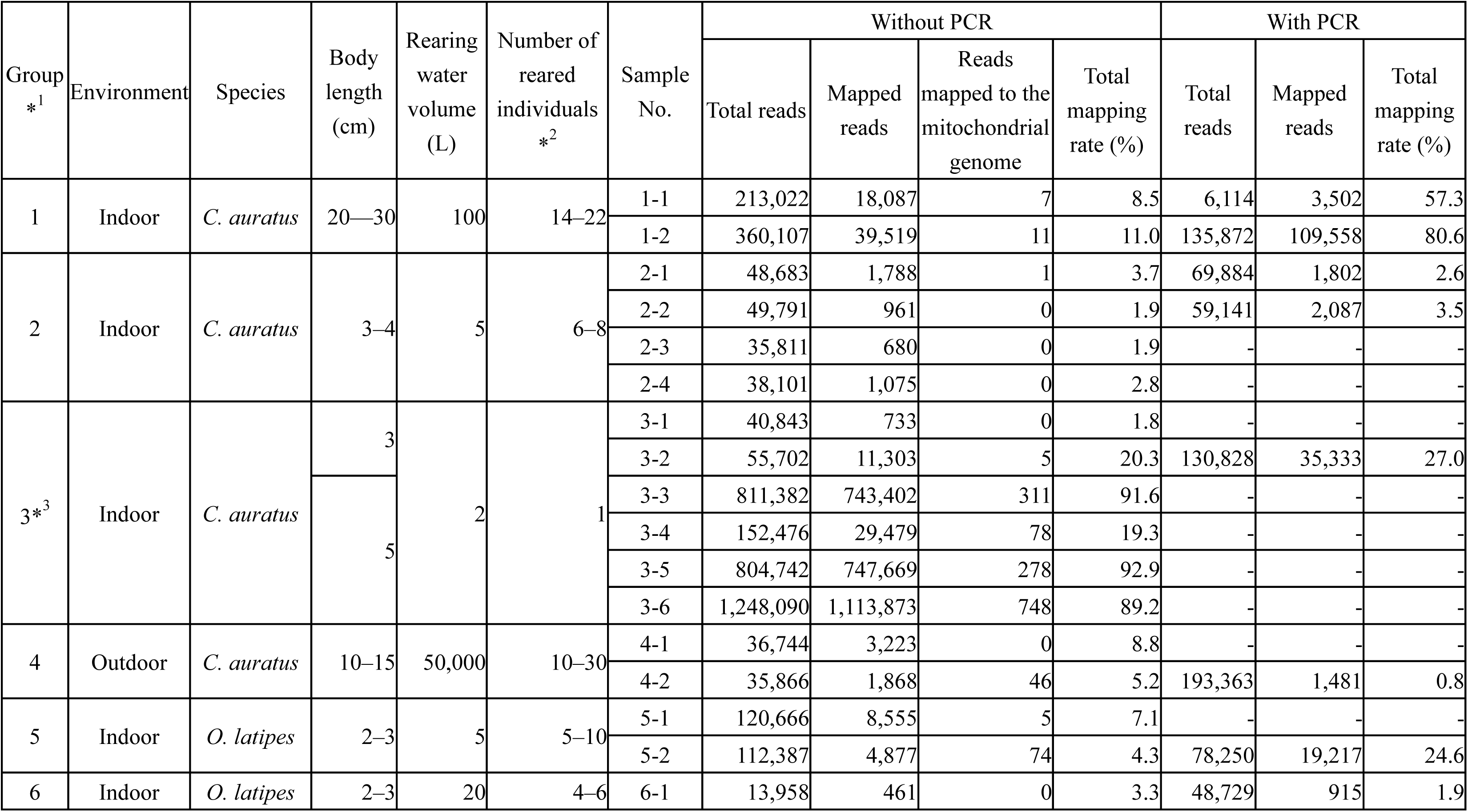
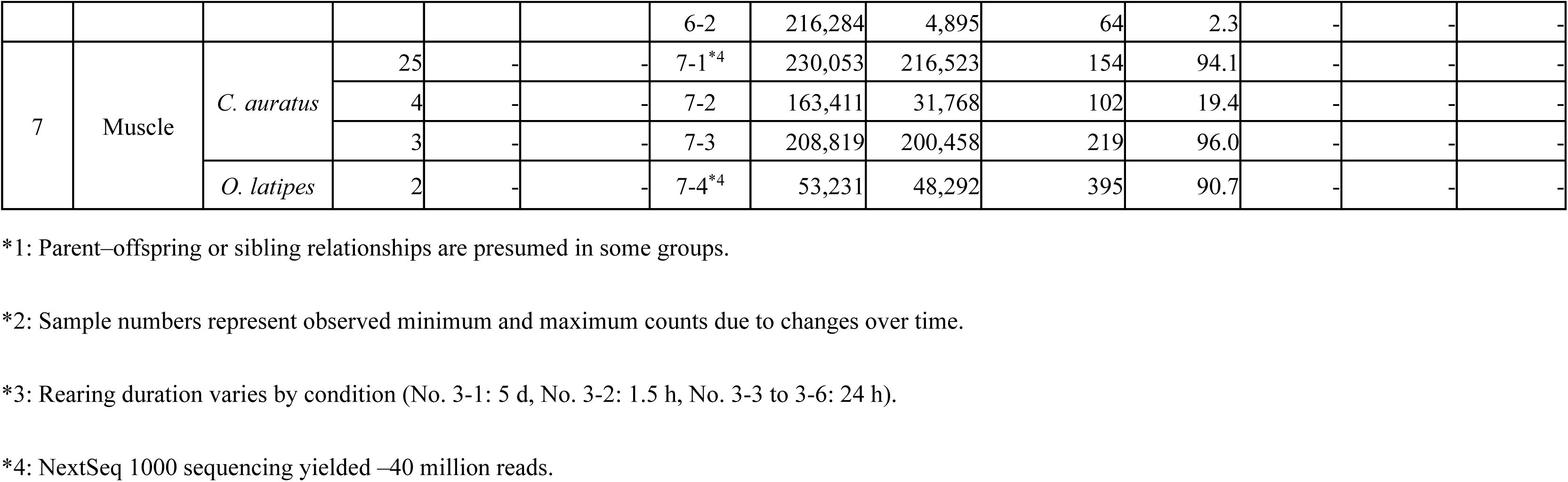
Rearing conditions for each tank, the collected samples, and their sequencing mapping results (with and without polymerase chain reaction [PCR])

#### 2.2.2. Long-range PCR of the entire mitochondrial genome

We amplified the full-length mitochondrial genome from aquarium water DNA as previously described (Deiner et al., 2017b). Each PCR mixture contained 25 µL of LongAmp® Taq 2× Master Mix (New England BioLabs, Ipswich, MA, USA), 15 µL of the aquarium-derived DNA extract diluted to 1.5 ng/µL, 5 µL of the forward primer Actinopterygii16SLRpcr_F (5′-CAGGA CATCC TAATG GTGCA G-3′) (4 pmol/µL), and 5 µL of the reverse primer Actinopterygii16SLRpcr_R (5′-ATCCA ACATC GAGGT CGTAA AC-3′) (4 pmol/µL). The thermal cycling protocol was as follows: initial denaturation at 94 °C for 30 s, followed by 35 cycles of denaturation (94 °C, 30 s), annealing (62 °C, 1 min), and extension (65 °C, 14 min 10 s), then a final extension at 65 °C for 10 min.

After PCR, the products were electrophoresed on a 0.8% agarose gel (1 h, 50 V). The band (16,000–17,000 bp) corresponding to full-length mitochondrial DNA was excised and purified using the QIAEX II Gel Extraction Kit (QIAGEN).

#### 2.2.3. Sequencing and bioinformatics

Libraries were prepared using the Ligation sequencing gDNA - Native Barcoding Kit 24 V14; SQK-NBD114.24 following the manufacturer’s instructions, from either the raw aquarium water/muscle tissue DNA extracts or the post-PCR DNA (Table 1). Sequencing was performed on a MinION Mk1C with an FLO-MIN114, R10.4.1 flow cell. Pod5 files were processed as described in Section 2.1.2 to generate fastq files. Reads were mapped using Minimap2 v2.28 to produce BAM files, which were visualised with SAMtools v1.21, IGV, and plot-bamstats. Short-read sequencing was performed on a NextSeq1000 (Illumina, San Diego, CA, USA) for one *C. auratus* (Table 1 No.7-1) and one *O. latipes* muscle tissue sample (Table 1 No.7-4). Libraries were prepared using the NEBNext® Ultra™ Ⅱ DNA PCR-free Library Prep Kit (New England Biolabs). After trimming and quality filtering with fastp v0.24.0 (Chen et al., 2018), reads were mapped using NextGenMap v0.5.5 (Sedlazeck et al., 2013). For both long- and short-read data, reference genomes were GCF_003368295.1/NC_006580.1 for *C. auratus* and GCF_002234675.1/NC_004387.1 for *O. latipes*.

*De novo* assembly was performed with Flye v2.9.4 (Lin et al., 2016; Kolmogorov et al., 2019) on long-read data from aquarium water (*C. auratus* and *O. latipes*: Table 1 Nos. 3-2 to 3-6, 5-2) and muscle tissue (Table 1 Nos. 7-1, 7-4). For some samples, reads were extracted or subsampled using SAMtools v1.21 and Trycycler v0.5.5 (Wick et al., 2021). Short-read mitochondrial reads (Table 1 Nos. 7-1, 7-4) were extracted with SAMtools v1.21, and the mitochondrial sequences were assembled using NOVOPlasty v4.3.5 (Dierckxsens et al., 2017). Final mitochondrial sequences were aligned in MEGA11 (Tamura et al., 2021), and variants identified manually or with SNP-sites v2.5.1 (Page et al., 2016).

To assess the benefits of complete mitochondrial sequencing, we downloaded full-length mitochondrial genomes of 12 *Carassius* species from MitoFish (Iwasaki et al., 2013). The sequences were aligned with MAFFT v7.525 (Katoh & Standley, 2013), and SNPs across full-length mitogenomes were identified using SNP sites v2.5.1 (Page et al., 2016) and counted with VCFtools v0.1.17 (Danecek et al., 2011). A 220-bp MiFish amplicon region (Miya et al., 2020) was extracted using Genetyx v15 (Software Development Co., Ltd., Toyama/Tokyo, Japan), aligned with Clustalw v1.83 (Thompson et al., 1994), and SNP positions and genotypes were visually determined.

### 2.3. Ethics statement

While muscle tissue was obtained from *C. auratus* and *O. latipes* individuals that died naturally (e.g., due to illness or accidental jumping), no animals were euthanised or subjected to invasive procedures. As such, ethical approval was not required under relevant Japanese regulations and institutional policies. All procedures were conducted in accordance with ASAB/ABS animal welfare guidelines.

## 3. Results

### 3.1. Effective experimental workflow for recovering long-fragment DNA of the target individuals from eDNA in water

This study aimed to establish an effective workflow for extracting long DNA fragments, including full-length mitochondrial genomes, from aquatic eDNA to maximise the benefits of LRS (Fig. 1, Consideration 1). We first extracted genomic DNA from aquarium water of *C. auratus* using four DNA extraction kits and compared the result, focusing on DNA yield and its length distribution (Fig. 2). The DNeasy Blood & Tissue Kit yielded the highest DNA concentration (4.11 ng/µL) but showed a peak fragment length of 22,683 bp, indicating the presence of many shorter fragments. The Extractor® WB Kit and MagAttract HMW DNA Kit both produced peaks >60,000 bp but also showed secondary peaks near 500 bp or had lower DNA yields. In contrast, the Blood & Cell Culture DNA Mini Kit generated a primary peak >60,000 bp without short-fragment peaks and had a relatively high DNA concentration (1.56 ng/µL), making it the most suitable for downstream applications. Based on these results, we selected this kit for subsequent experiments.

**Figure 2.**
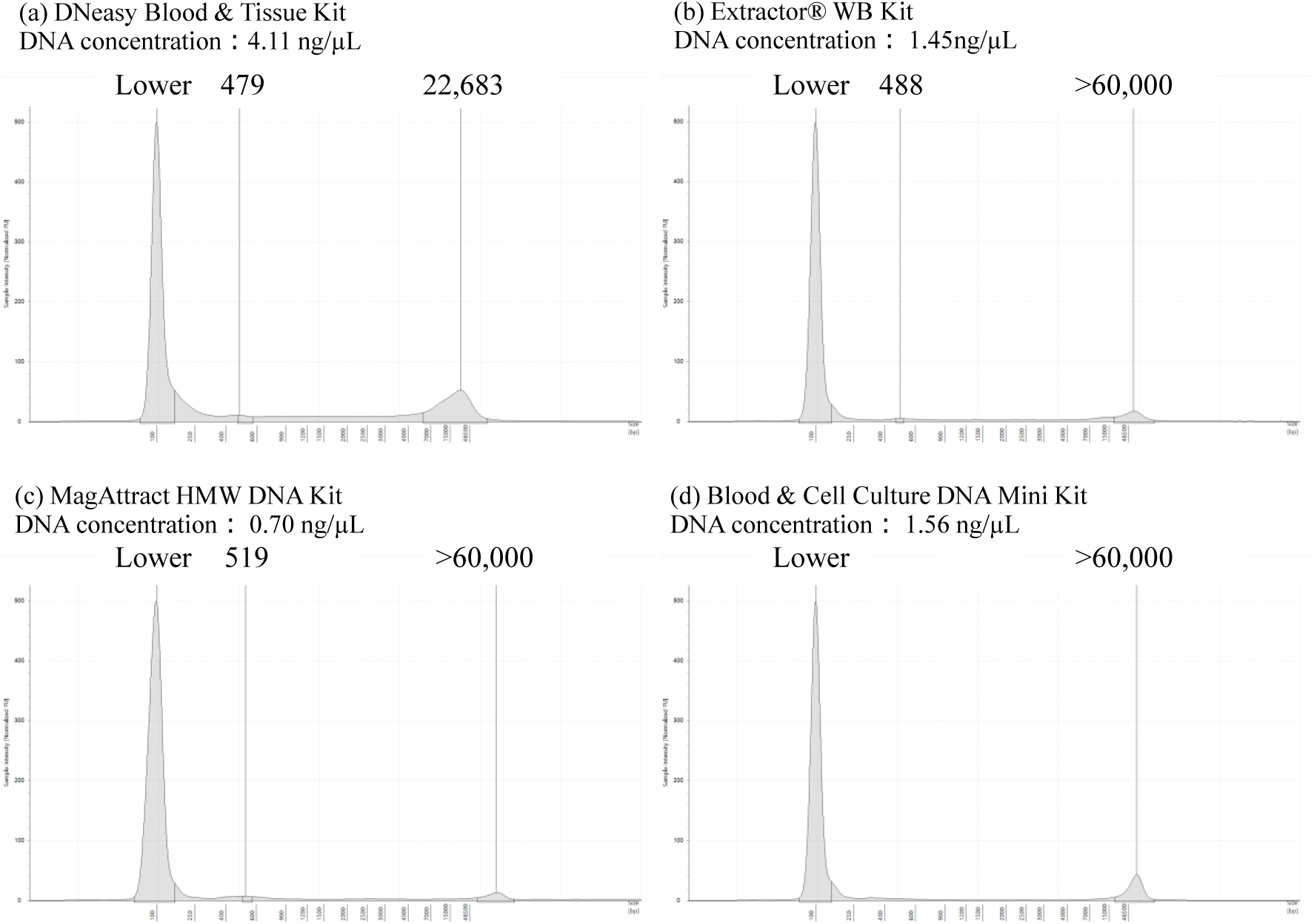
Fragment length distribution measured by TapeStation and DNA concentration measured by Qubit® 2.0 Fluorometer for extracts obtained using different extraction kits. (a) DNeasy Blood & Tissue Kit, (b) Extractor® WB Kit, (c) MagAttract HMW DNA Kit, (d) Blood & Cell Culture DNA Mini Kit. Peaks corresponding to the maximum measurable fragment length (>60,000 bp) were observed in all kits except (a).

We then assessed membrane filter pore sizes using water from aquaria containing *C. auratus* and *O. latipes*. All tested filters yielded primary peaks >60,000 bp with no major differences in fragment length (Supplementary Fig. S1). Among them, the 0.2 µm filter yielded the highest DNA concentration (17 ng/µL), while other pore sizes ranged from 0.35 to 2.8 ng/µL. To evaluate target sequence recovery, we compared mapping rates against whole reference genomes. Filters with 1.2 µm pores achieved the highest mapping rates for both species, while larger or smaller pores resulted in reduced recovery (Fig. 3; Supplementary Table S1).

**Figure 3.**
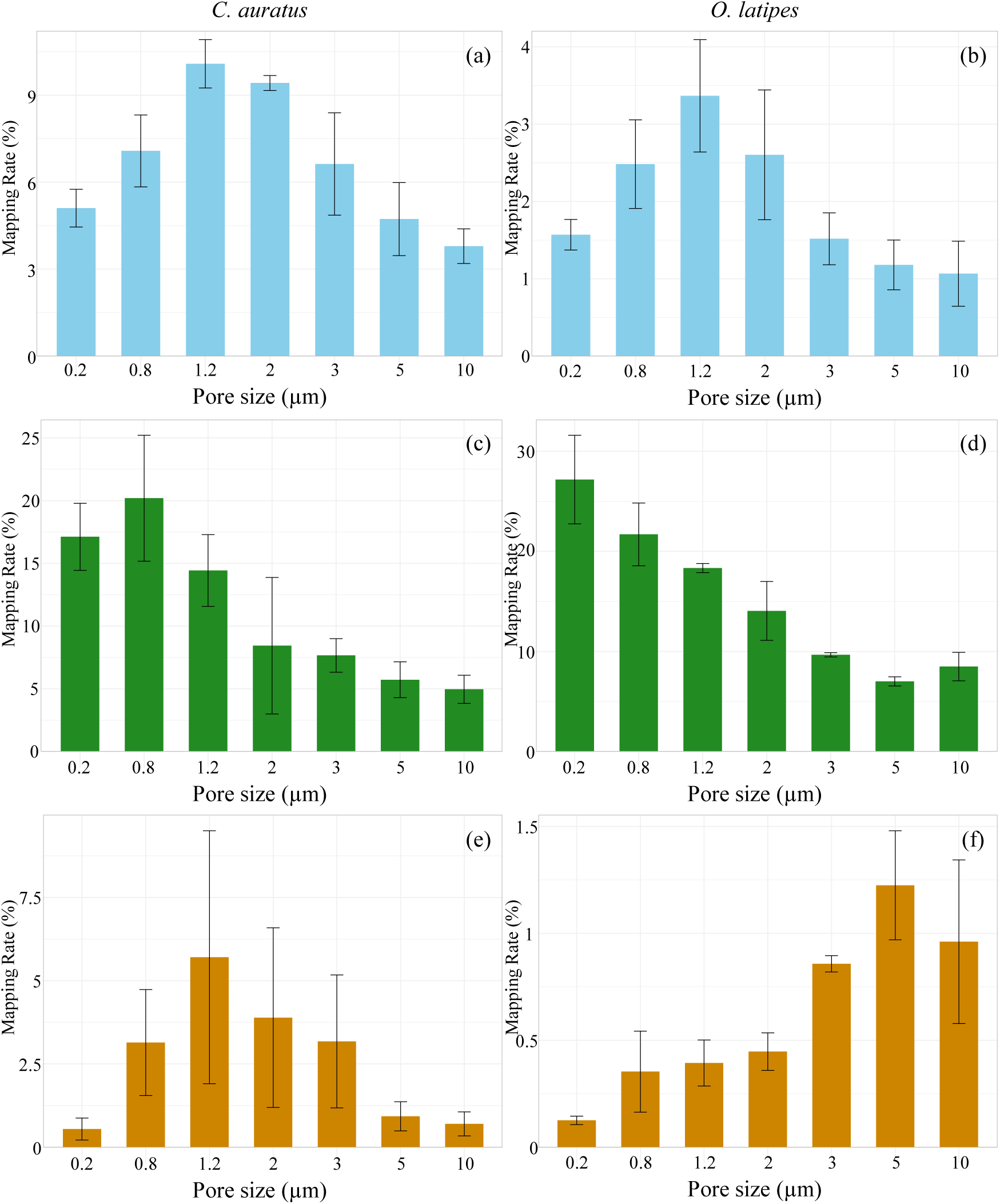
Mapping rates and proportions of bacterial/fungal DNA by filter pore size (a) *C. auratus*: Mapping rate, (b) *O. latipes*: Mapping rate, (c) *C. auratus*: Bacterial DNA, (d) *O. latipes*: Bacterial DNA, (e) *C. auratus*: Fungal DNA, (f) *O. latipes*: Fungal DNA Graphs show average values from three replicates with standard deviations.

To assess the impact of non-target DNA, we also analysed bacterial and fungal DNA proportions. Bacterial DNA was more abundant at smaller pore sizes (up to 27.2%), whereas fungal DNA generally remained below 10%, showing species- and pore size-specific trends (Fig. 3).

### 3.2. Sequence data acquisition from different water samples

After selecting an optimal extraction method and filter pore size (Blood & Cell Culture DNA Mini Kit with a 1.2 µm filter), we applied two sequencing strategies: a PCR-free approach to obtain complete mitochondrial genomes without amplification bias (Fig. 1, Consideration 2), and a PCR-based approach using published primers for whole-mitogenome amplification to recover sequences from low-yield samples (Fig. 1, Consideration 3).

#### 3.2.1 PCR-free analysis

We conducted LRS of aquatic eDNA without PCR amplification. The mapping rates against the whole-genome sequence varied widely, ranging from a few percent to nearly 90%, depending on rearing conditions (Table 1). Factors including indoor vs. outdoor environment, rearing density, sampling day, and rearing duration influenced the results. For instance, comparing No.3-1 (5 d rearing) and No.3-2 (1.5 h rearing), the latter showed a higher mapping rate (Table 1). In some single-fish tanks of *C. auratus* (Table 1, Nos. 3-3, 3-5, 3-6), aquarium-derived DNA yielded tens to hundreds of reads mapped to the mitochondrial genome, some of which spanned nearly the full length. For example, 748 reads were obtained in No. 3-6 with approximately uniform coverage (Supplementary Fig. S2). Among these, 17 reads were >16,000 bp. A homology search using blastn (Altschul et al., 1990) identified the top hit as the *C. auratus* mitochondrial genome (MF443764.1), strongly suggesting that these reads originated from *C. auratus* mitochondria.

#### 3.2.2 PCR-based analysis

We performed PCR amplification of the full-length Actinopterygii mitochondrial genome for eDNA from aquarium water under various conditions, followed by LRS. Enrichment efficiency varied by sample. In several samples, more mitochondria-mapped reads were obtained compared to PCR-free sequencing (Table 1). Notably, mitochondrial reads in sample No. 1-2 increased from 11 (without PCR) to 109,450 (with amplification). The average read length was 9,013 bases, with 27.4% exceeding 16,000 bases—sufficient to cover the complete mitochondrial genome. Coverage was 100% with a mean depth of 58,434.1, and alignment visualisation revealed uniform coverage except near the primer regions (3,436–3,478 bp) (Supplementary Fig. S3). Conversely, in sample No. 4-2, only 0.8% of reads mapped to the mitochondrial genome even after PCR.

### 3.3. *De novo* assembly of the full-length mitochondrial genome

We performed *de novo* assembly of full mitochondrial genomes using eDNA and muscle tissue-derived DNA (Fig. 1, Consideration 2 and 3), including PCR-free aquarium samples from individually reared *C. auratus* (Table 1 Nos. 3-3 to 3-6) and PCR-amplified aquarium water from *C. auratus* or *O. latipes* (Nos. 3-2 and 5-2). In addition, muscle tissue samples (Nos. 7-1 and 7-4) were included as invasive controls to provide ground-truth mitochondrial sequences for comparison. As part of the sample selection criteria, all aquarium water samples—except those from *O. latipes*—contained a single fish. Muscle samples were also sequenced with NextSeq 1000 for comparison with short-read assemblies.

First, we evaluated assembly accuracy by comparing short- and long-read assemblies from muscle-derived DNA. In *C. auratus*, both assemblies were identical, confirming the accuracy of long-read assembly. In contrast, the short-read assembly of *O. latipes* did not yield a complete mitochondrial genome, resulting in a pairwise identity of 99.95 %.

We defined a successful assembly as a single contiguous sequence covering the full mitochondrial genome (i.e., 16,580 bp for *C. auratus*, 16,714 bp for *O. latipes*) with ≥99.9% identity to a species-specific reference in the NCBI database, as determined by BLASTn (Altschul et al., 1990). Based on this criterion, we first evaluated PCR-free water samples from individually reared *C. auratus* (Table 1, Nos. 3-3 to 3-6). In these cases, assemblies were consistently successful when only mitochondria-mapped reads were used as input, demonstrating that high-quality mitochondrial genomes can be reconstructed from eDNA without amplification. We then assessed samples obtained via PCR amplification of aquarium water and muscle tissue. When raw reads were directly input into the assembler, assemblies sometimes failed, likely due to read length variation or sequencing errors. However, by optimising preprocessing procedures—specifically extracting reads mapped to the mitochondrial genome, filtering by read length (16,000– 18,000 bp), and subsampling to remove excess reads—successful assemblies were achieved for all tested samples (Table 2). In particular, assemblies tended to be more successful with fewer input reads.

**Table 2.**
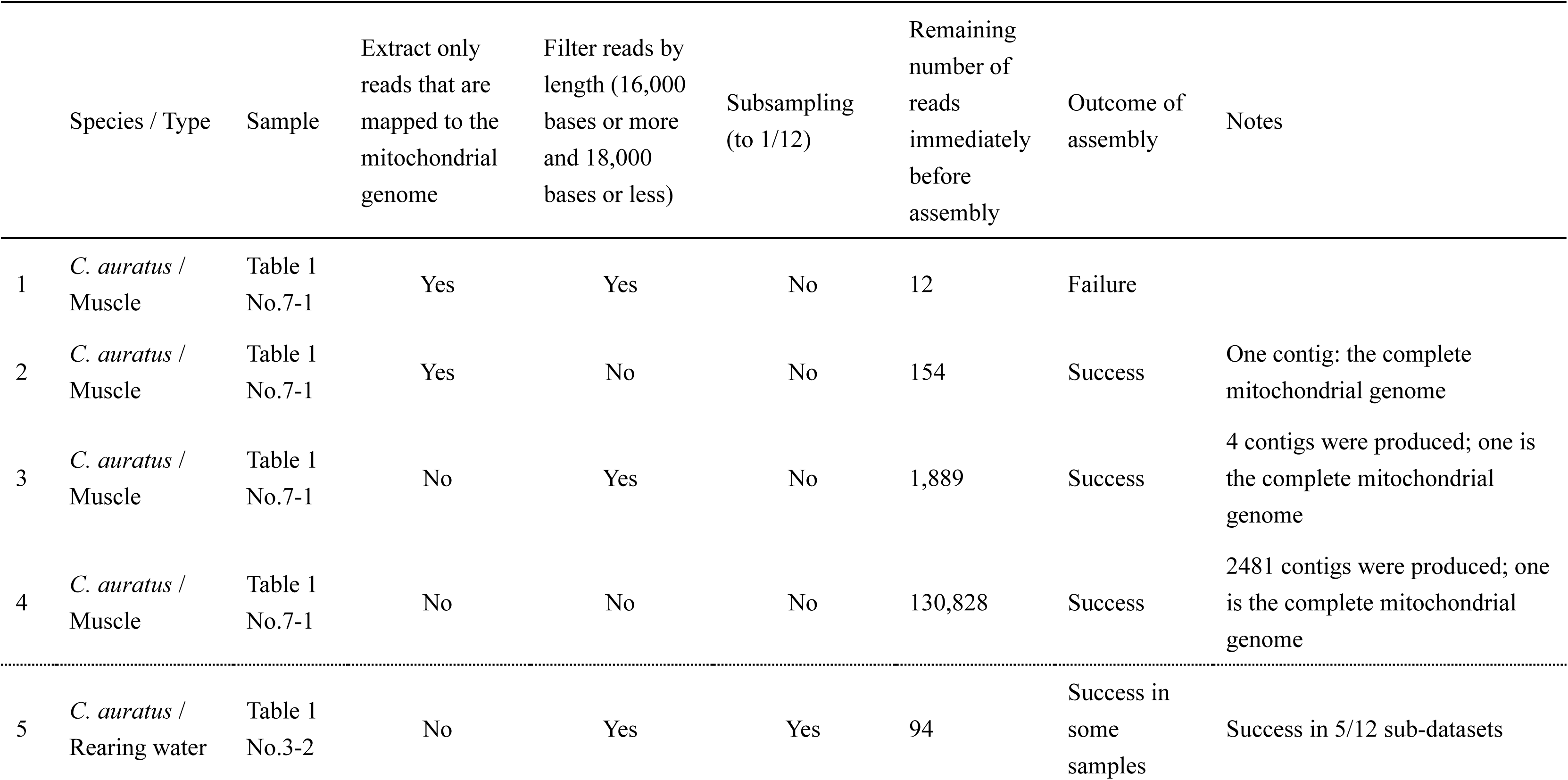
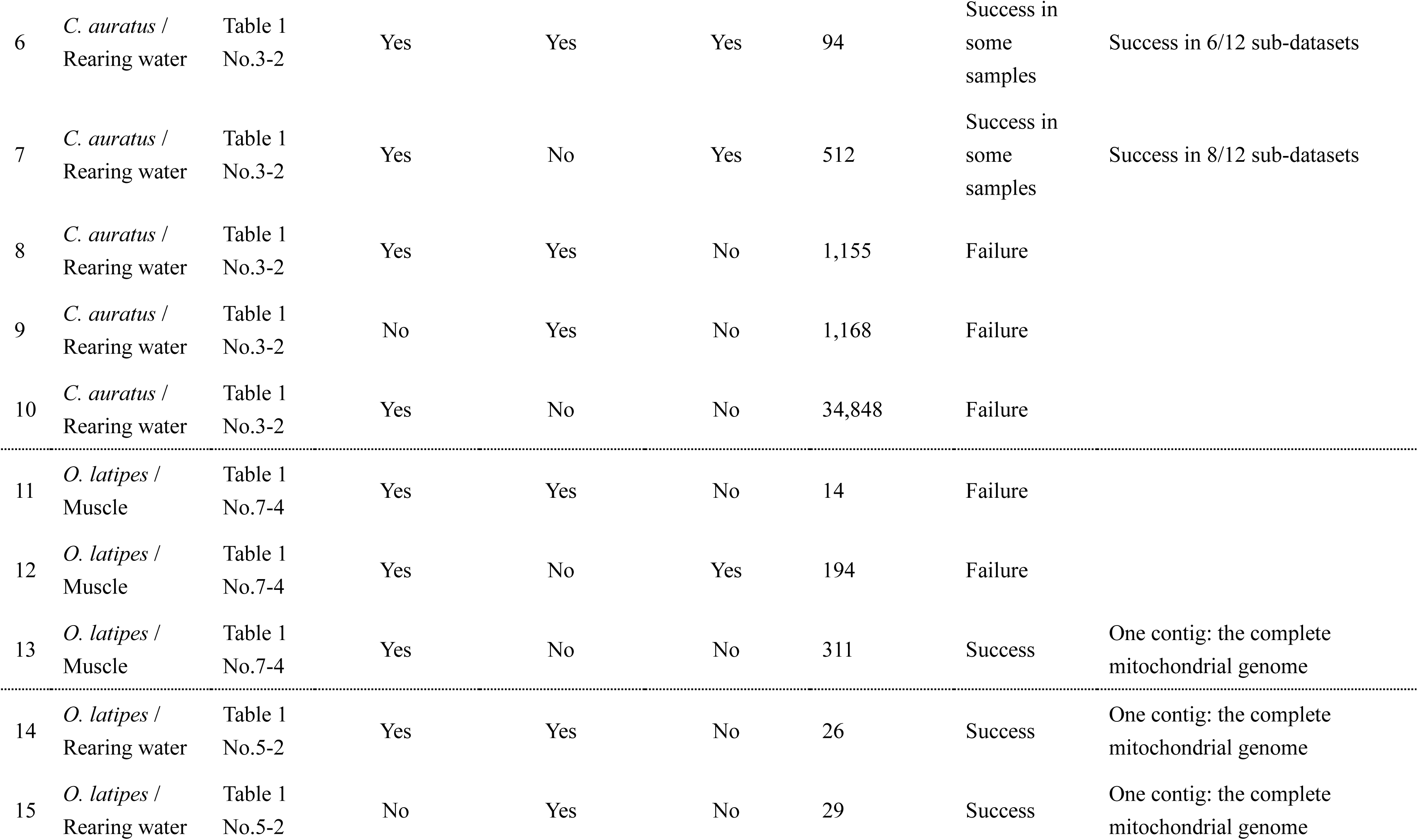
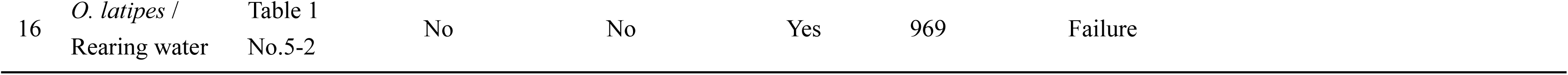
Summary of workflows and outcomes for *de novo* assembly of mitochondrial genomes.

|  | Species / Type | Sample | Extract only<br>reads that are<br>mapped to the<br>mitochondrial<br>genome | Filter reads by<br>length (16,000<br>bases or more<br>and 18,000<br>bases or less) | Subsampling<br>(to 1/12) | Remaining<br>number of<br>reads<br>immediately<br>before<br>assembly | Outcome of<br>assembly | Notes |
| --- | --- | --- | --- | --- | --- | --- | --- | --- |
| 1 | *C. auratus* /<br>Muscle | Table 1<br>No.7-1 | Yes | Yes | No | 12 | Failure |  |
| 2 | *C. auratus* /<br>Muscle | Table 1<br>No.7-1 | Yes | No | No | 154 | Success | One contig: the complete<br>mitochondrial genome |
| 3 | *C. auratus* /<br>Muscle | Table 1<br>No.7-1 | No | Yes | No | 1,889 | Success | 4 contigs were produced; one is<br>the complete mitochondrial<br>genome |
| 4 | *C. auratus* /<br>Muscle | Table 1<br>No.7-1 | No | No | No | 130,828 | Success | 2481 contigs were produced; one<br>is the complete mitochondrial<br>genome |
| 5 | *C. auratus* /<br>Rearing water | Table 1<br>No.3-2 | No | Yes | Yes | 94 | Success in<br>some<br>samples | Success in 5/12 sub-datasets |
| 6 | *C. auratus* /<br>Rearing water | Table 1<br>No.3-2 | Yes | Yes | Yes | 94 | Success in<br>some<br>samples | Success in 6/12 sub-datasets |
| 7 | *C. auratus* /<br>Rearing water | Table 1<br>No.3-2 | Yes | No | Yes | 512 | Success in<br>some<br>samples | Success in 8/12 sub-datasets |
| 8 | *C. auratus* /<br>Rearing water | Table 1<br>No.3-2 | Yes | Yes | No | 1,155 | Failure |  |
| 9 | *C. auratus* /<br>Rearing water | Table 1<br>No.3-2 | No | Yes | No | 1,168 | Failure |  |
| 10 | *C. auratus* /<br>Rearing water | Table 1<br>No.3-2 | Yes | No | No | 34,848 | Failure |  |
| 11 | *O. latipes* /<br>Muscle | Table 1<br>No.7-4 | Yes | Yes | No | 14 | Failure |  |
| 12 | *O. latipes* /<br>Muscle | Table 1<br>No.7-4 | Yes | No | Yes | 194 | Failure |  |
| 13 | *O. latipes* /<br>Muscle | Table 1<br>No.7-4 | Yes | No | No | 311 | Success | One contig: the complete<br>mitochondrial genome |
| 14 | *O. latipes* /<br>Rearing water | Table 1<br>No.5-2 | Yes | Yes | No | 26 | Success | One contig: the complete<br>mitochondrial genome |
| 15 | *O. latipes* /<br>Rearing water | Table 1<br>No.5-2 | No | Yes | No | 29 | Success | One contig: the complete<br>mitochondrial genome |

|  |  |  |  |  |  |  |  |
| --- | --- | --- | --- | --- | --- | --- | --- |
| 16 | *O. latipes* /<br>Rearing water | Table 1<br>No.5-2 | No | No | Yes | 969 | Failure |

### 3.4. Polymorphism analysis across the full-length mitochondrial genome

We next aligned the assembled mitochondrial genomes and analysed single nucleotide variants (SNVs) (Fig. 1 Consideration 4). For *C. auratus*, six samples (aquarium water and muscle tissue) shared two SNV sites across the mitochondrial genome. Three samples—PCR-amplified water (Table 1 No. 3-2), PCR-free water (No. 3-6), and muscle (No. 7-1)—showed identical sequences, while the remaining three PCR-free water samples (Nos. 3-3, 3-4, 3-5) also matched each other (Figs. 4 and 5). These results indicated two haplotypes distinguished by two SNVs: a single-nucleotide substitution in the D-loop (site 1) and a deletion in the 16S rRNA region (site 2) (Supplementary Table S2). Read distribution at these sites, visualised using IGV, showed that for site 1, approximately 98% of reads in samples No. 3-2 and No. 7-1 supported the substitution, while in No. 3-6, 64% supported the substitution and about 34% matched the reference (Fig. 6A, Supplementary Fig. S4). For site 2, assemblies of samples Nos. 3-2, 3-6, and 7-1 contained the deletion; in No. 3-6, ∼65% of reads carried the deletion (Fig. 6B, Supplementary Fig. S5). In No. 3-6, reads >16,000 bp spanning both SNVs suggested the presence of two haplotypes (Supplementary Figs. S6 and S7). Additionally, comparison of PCR-amplified water sample No. 3-2 with the reference genome revealed 44 SNVs.

**Figure 4.**
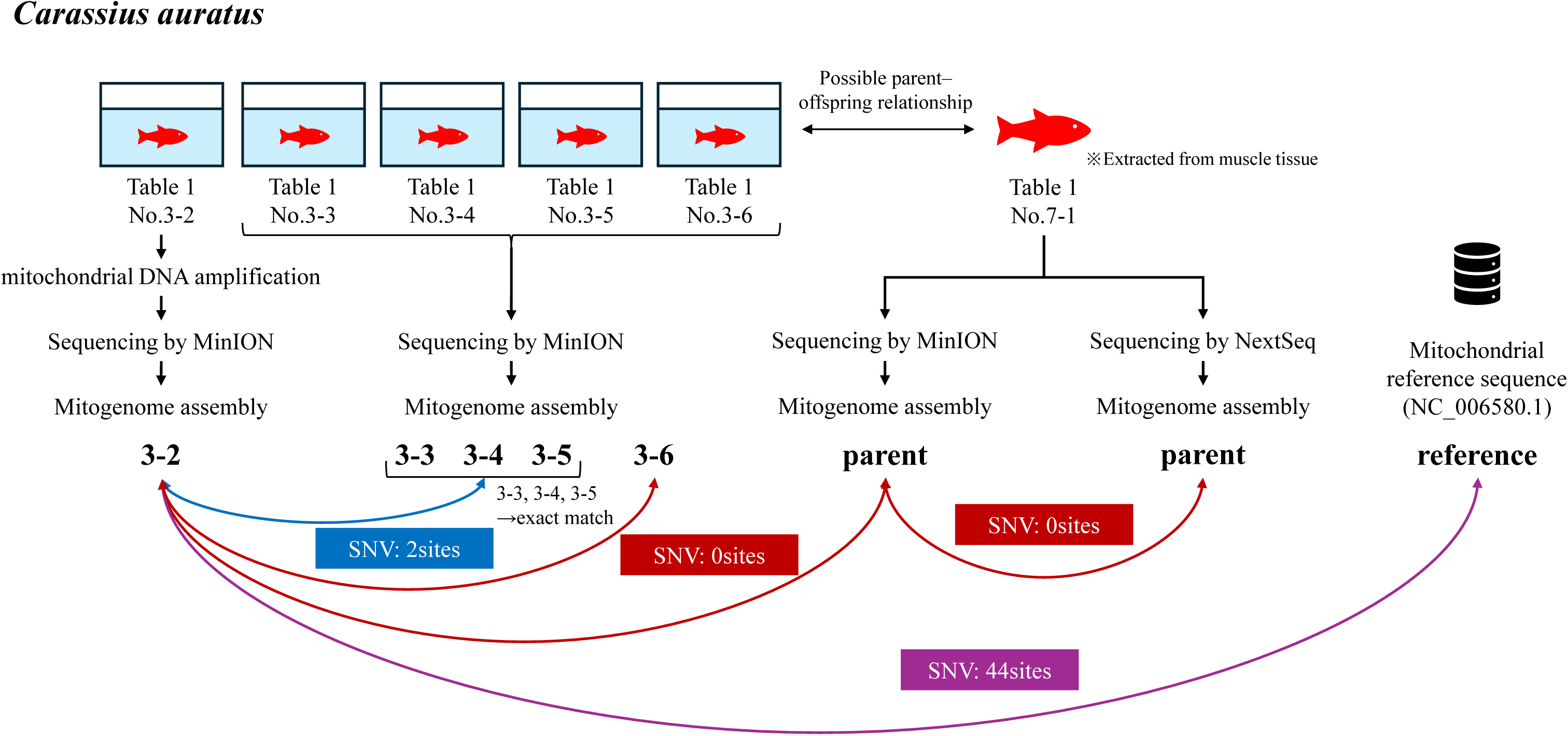
Workflow of *de novo* mitochondrial assembly and polymorphism analysis (*C. auratus*) Comparison of mitochondrial genome assemblies from single-fish tanks and a muscle sample. Two haplotype groups differing at two SNV sites were identified No.3-2 differed from the reference at 44 SNV sites.

**Figure 5.**
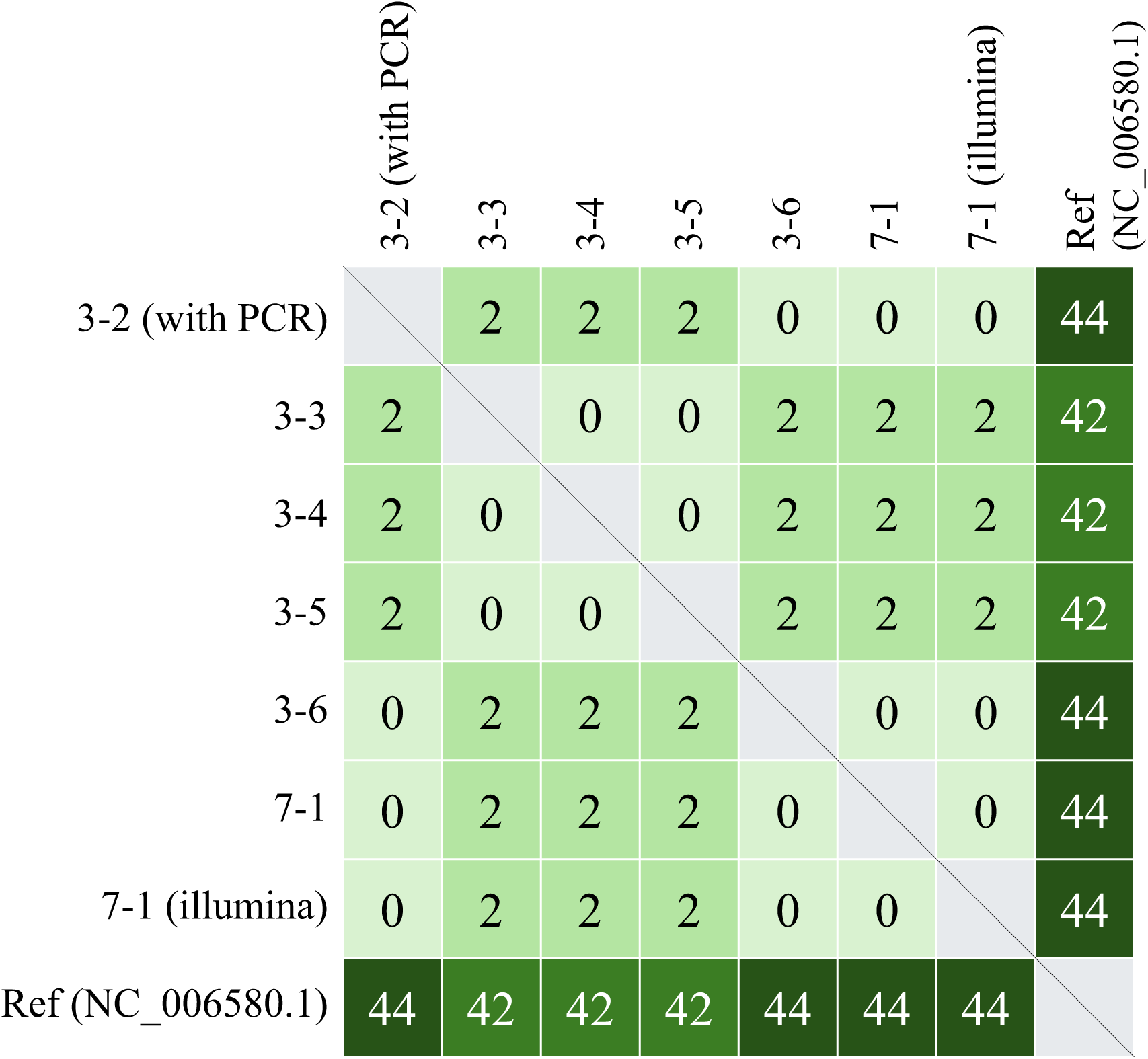
Alignment of full-length mitochondrial genomes among *C. auratus* samples Each number indicates the count of SNVs detected across the full mitochondrial genome in pairwise comparisons between samples.

**Figure 6.**
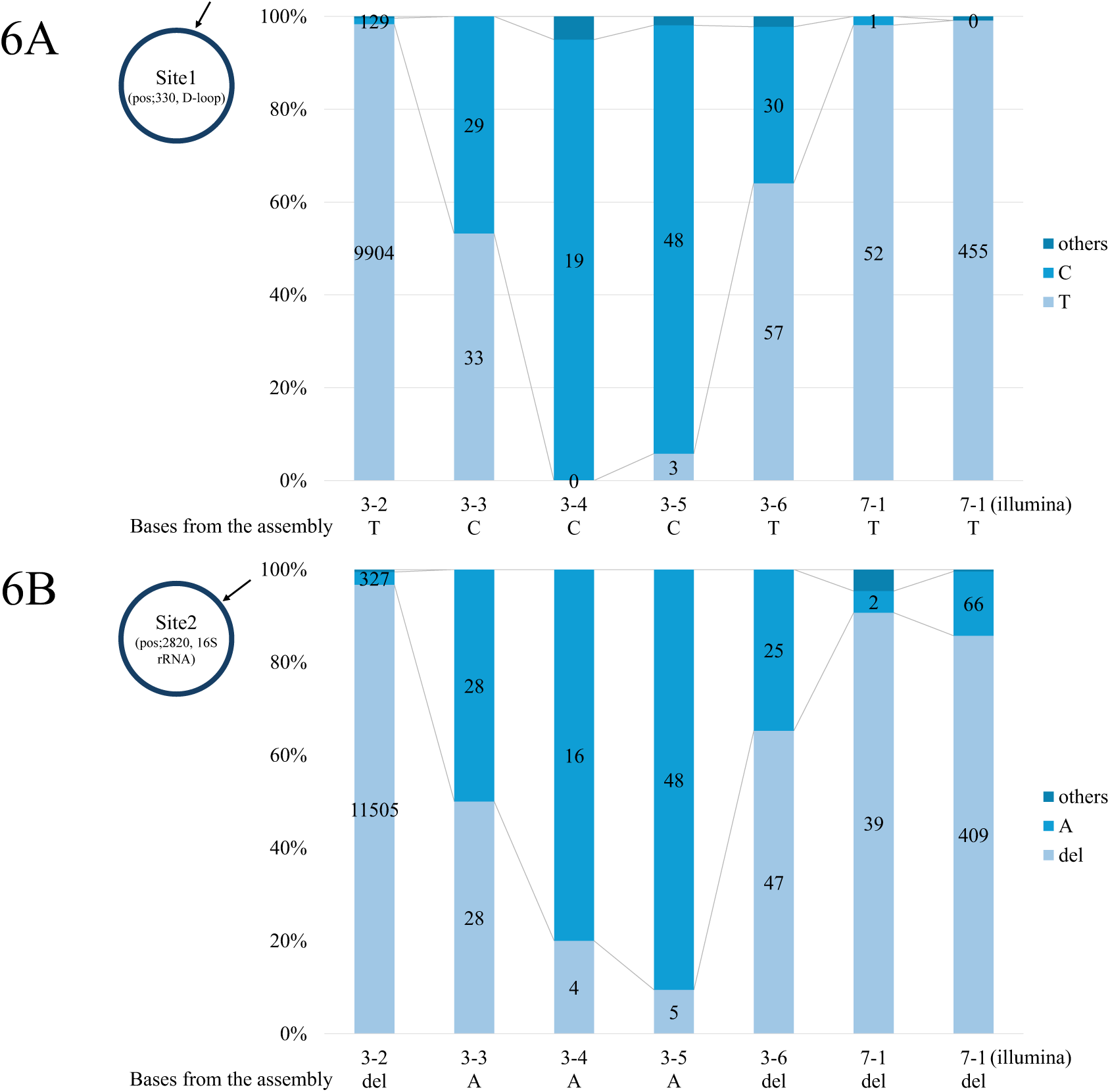
Read distributions at two SNV sites in the mitochondrial genome of *C. auratus* (A) Read counts for each allele (C, T, or others) at SNV site 1 located in the D-loop region. (B) Read counts for each allele (A, deletion, or others) at SNV site 2 located in the 16S rRNA region. For both panels, allele-specific read counts are shown for each sample, with the assembled allele indicated below each plot.

For *O. latipes*, comparison between the PCR-amplified aquarium water sample (Table 1 No. 5-2) and the muscle sample (No. 7-4) identified two SNVs, and 314 SNVs were detected relative to the reference genome (Fig. 7).

**Figure 7.**
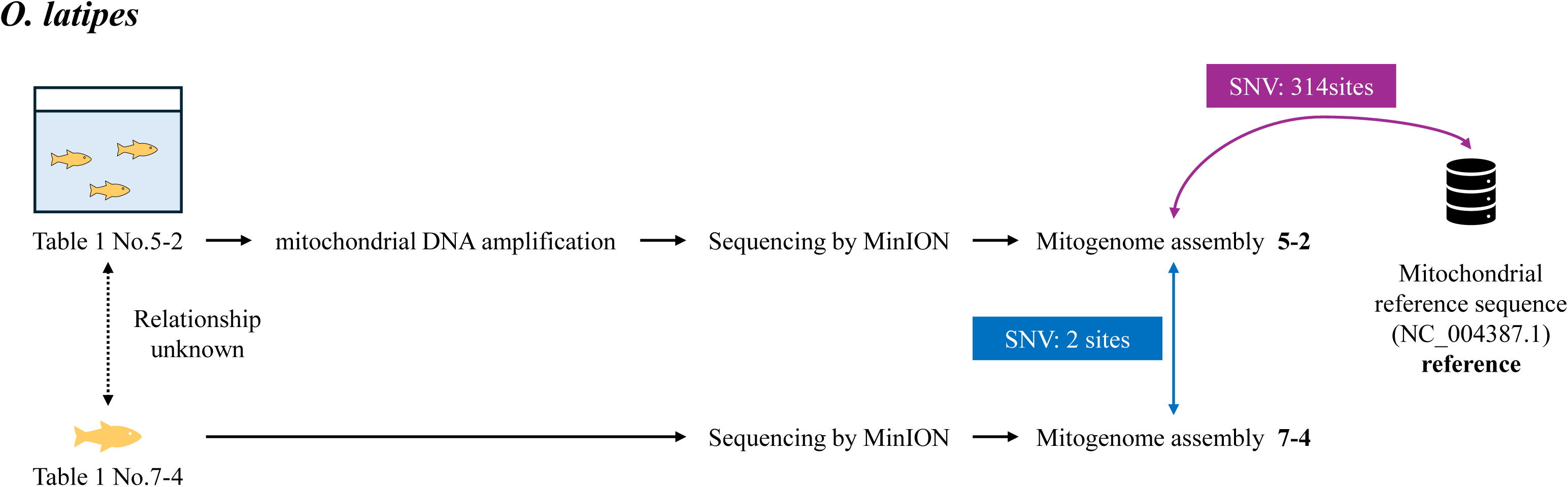
Workflow of *de novo* mitochondrial assembly and SNV analysis for *O. latipes* A comparison of PCR-amplified aquarium water samples and muscle tissue revealed 2 SNVs and 314 SNVs relative to the reference.

### 3.5. Advantage of sequencing the complete *Carassius* mitochondrial genome

To assess the value of sequencing the complete mitochondrial genome, we compared SNV counts and genotypes between full-length and partial mitochondrial sequences. Table 3 summarises SNV counts in the full mitochondrial genomes and the 220-bp MiFish region of 12 Carassius species. In total, 2,415 SNVs were identified in the full-length sequences. SNVs were identified relative to a pseudo-reference generated from the multiple-sequence alignment. Several species had approximately 600–700 SNVs, with the highest in the twelfth sample, a hybrid between *Cyprinus carpio wuyuanensis* and *C. auratus.* In contrast, only five SNVs were detected within the MiFish region among the 12 species. As shown in Table 3, species 1–5 (*C. langsdorfii* to *C. cuvieri*) shared identical genotypes, as did species 9 (*C. gibelio*) and 10 (*C. auratus* ssp. Pingxiang), making them indistinguishable based on MiFish markers.

**Table 3.** Polymorphism counts in mitochondrial genomes of 12 *Carassius* species.

|  | Species name | Number of single nucleotide variants (SNVs) in the whole mitochondrial genome | The nucleotides of SNVs in the 220-bp region amplified by MiFish primers |  |  |  |  |
| --- | --- | --- | --- | --- | --- | --- | --- |
| 1 | *C. langsdorfii* | 3 | T | C | G | A | G |
| 2 | *C. auratus* × *Cyprinus carpio* | 649 | T | C | G | A | G |
| 3 | *C. auratus* × *Cyprinus carpio* × *C. cuvieri* | 671 | T | C | G | A | G |
| 4 | *C. auratus auratus* | 664 | T | C | G | A | G |
| 5 | *C. cuvieri* | 761 | T | C | G | A | G |
| 6 | *C. auratus* × *Megalobrama amblycephala* tetraploid hybrid | 641 | T | C | A | A | G |
| 7 | *C. auratus* × *Megalobrama amblycephala* triploid hybrid | 648 | T | C | A | A | G |
| 8 | *C. auratus* × *Megalobrama amblycephala* pentaploid hybrid | 755 | T | T | A | G | G |
| 9 | *C. gibelio* | 651 | C | C | A | G | A |
| 10 | *C. auratus* ssp Pingxiang | 663 | C | C | A | G | A |
| 11 | *Cyprinus carpio wuyuanensis* × *C. auratus* | 1,641 | C | C | A | A | A |
| 12 | *C. auratus* × *Megalobrama amblycephala* × *C. cuvieri* | 680 | C | A | A | A | A |
| All |  | 2,415 |  |  |  |  |  |

## 4. Discussion

Herein, we developed an eDNA-specific filtration workflow coupled with PCR-free LRS (Fig. 1), we recovered complete mitochondrial genomes from as little as 250 mL of aquarium water, yielding ≥99.9 % identity to reference sequences. This technical advance has the promise to move eDNA applications beyond species detection toward population-level genomics, setting the stage for assessing intraspecific diversity in natural systems.

To clarify how each experimental step contributed to this advance, we structure the discussion around four key components that mirror Figs 2–8. First, we optimised DNA extraction using a 1.2 µm pore-size membrane filter and a gravity-based extraction kit (Figs. 2 and 3), which minimises shearing and is suitable for high-molecular-weight DNA (Angthong et al., 2020). These trends are consistent with previous reports that smaller pores clog easily while larger ones cause particle loss (Liu et al., 2024). Mapping rates peaked at 1.2 µm and declined with both smaller and larger pores, likely due to increased non-target DNA (Thomsen & Willerslev, 2015). Smaller pores trapped more bacterial DNA, while larger pores retained more fungal DNA in *O. latipes*. The increase in fungal DNA was not observed in *C. auratus*, possibly due to increased loss of *C. auratus* DNA through the filter or algal contamination. Although 1.2 µm was optimal here, other studies found 0.2 µm (Turner et al., 2014) or 3.0 µm (Liu et al., 2024) to be more effective, suggesting that the optimal pore size depends on the research objective, downstream analyses, and criteria for evaluating performance. Additionally, mapping rates varied across samples despite identical methods, indicating that fish activity and sampling timing affect eDNA recovery (Klymus et al., 2015; Takahara et al., 2012).

**Figure 8.**
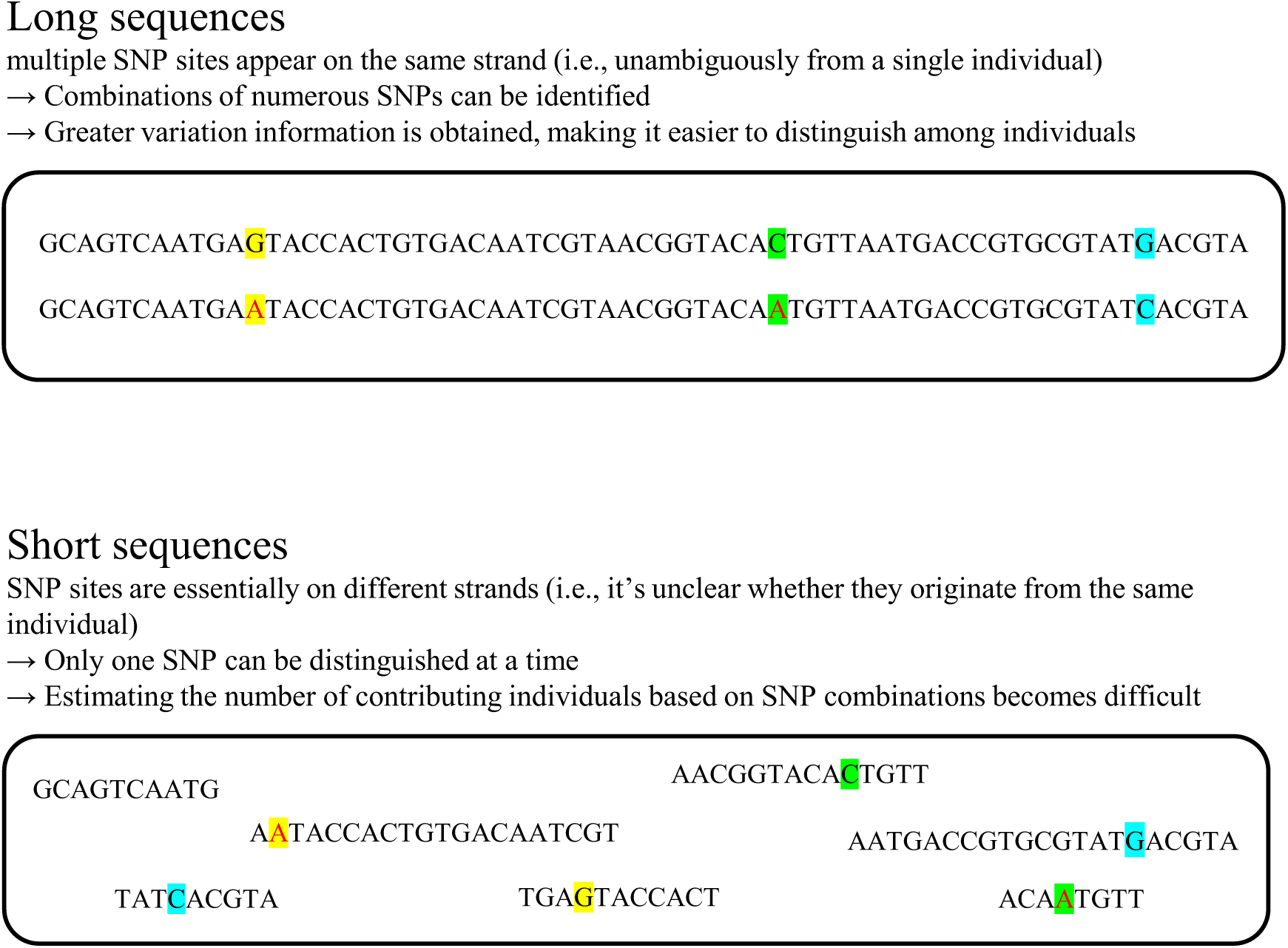
Future Idea: Advantage of long-read sequencing for polymorphism analysis Diagram illustrating how long-read sequencing enables multiple SNPs to be analysed on a single strand, improving the ability to infer individual haplotypes and increasing resolution compared to short-read methods.

We also demonstrated that full-length mitochondrial sequences can be recovered without PCR (Table 1 and Supplementary Fig. S2). Several PCR-free samples yielded near-full-length mitochondrial reads, suggesting the presence of intact mitochondria or mitochondrial DNA in aquatic environments (Karin et al., 2023; Liu et al., 2024). This approach avoids fragmentation, enabling detection of polymorphic sites on single molecules (Fig. 8) and improving genotype resolution in mixed populations (Sethi et al., 2019). Notably, while most PCR-free samples had low mapping rates (ranging from few per cent to about ten per cent), certain *C. auratus* samples (Table 1, Nos. 3-3, 3-5, 3-6) approached 90%, likely due to prompt sampling and high DNA yield from favourable body size-to-volume ratios (Dejean et al., 2011; Thomsen et al., 2012).

PCR amplification and LRS provided high-depth coverage across full mitochondrial genomes, far exceeding typical ∼220 bp eDNA barcodes. This enhances species discrimination in taxa that are indistinguishable with short markers (Dziedzic et al., 2023), facilitating noninvasive sampling in ethically or logistically constrained species. For instance, *Carassius* species indistinguishable by 220 bp sequences exhibited numerous polymorphisms in full-length data (Table 3), enhancing taxonomic precision. However, PCR efficiency varied; for example, the field sample (No. 4-1) contained mitochondrial DNA but showed poor amplification, likely due to inhibitors from the turbid water (Jane et al., 2015). Additionally, although not problematic in this study, chimeric reads should be considered with caution in downstream analyses.

We demonstrated that mitochondrial genomes can be assembled *de novo* from eDNA. Assembly success depended on species and PCR status (Table 2). For PCR-amplified samples, subsampling was required to avoid excess reads, which can hinder assembly (Deiner et al., 2017b). Accurate assemblies from water samples (Table 1, Nos. 3-2, 3-6) matched those from muscle tissue (No. 7-1), confirming the sequence quality. PCR-free assembly also succeeded in samples with high mitochondrial read abundance (Nos. 3-3 to 3-6). IGV visualisation showed multiple alleles at some polymorphic sites (Figs. 6A and B); in No. 3-6, two haplotypes differing by two SNVs appeared to coexist (Supplementary Figs. S6 and S7), possibly indicating heteroplasmy. Taken together, our results suggest that high-quality, individual-level mitochondrial genomes can be obtained from eDNA without species-specific primers.

As noninvasive sampling becomes increasingly important for conservation-priority species (Kakehashi et al., 2022), our framework offers a practical tool for assessing intraspecific diversity, informing taxonomy and phylogeny, and supporting genomic conservation (Ouborg et al., 2010). Interestingly, some PCR-free *C. auratus* samples showed exceptionally high mapping rates, likely due to high DNA release and prompt sampling, suggesting that this strategy is particularly effective under conditions with high eDNA concentrations and minimal degradation. While promising, this approach has limitations. First, we used few replicates due to a lack of aquarium space. Second, our studied did not control for environmental factors (e.g., turbidity, microbial composition, fish behaviour). Finally, we focused on mitochondrial rather than nuclear DNA, which may provide limited information for species with low mitochondrial diversity. Long-range PCR of nuclear regions may support more detailed analyses of population structure and adaptive variation (Krehenwinkel et al., 2019). However, applying this method to nuclear loci poses additional challenges, such as their lower copy numbers and greater fragmentation. Addressing these limitations—together with increased replication and broader sampling—will be essential for wider applications.

## Author contributions

Hinano Mizuno conceived the ideas, designed methodology, collected the data, analysed the data, and led the writing of the manuscript; Hidenori Tanaka, Yoshikazu Furuta, and Nobuhiko Muramoto revised the manuscript critically for important intellectual content. All authors contributed critically to the drafts and gave final approval for publication.

