## Supplementary Figures for "Noninvasive reconstruction of complete mitochondrial genomes from aquatic environmental DNA using PCR-free long-read sequencing"

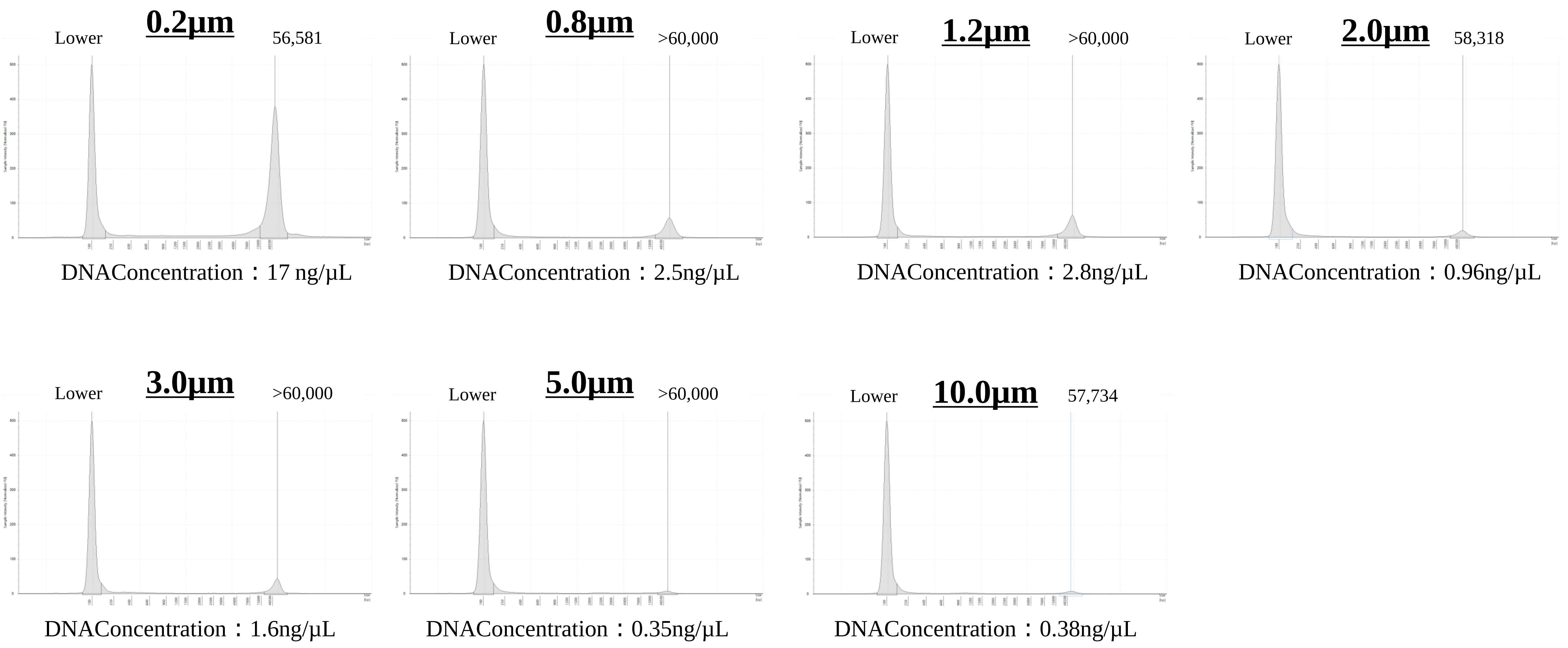


**Supplementary Figure S1. DNA fragment length distribution by filter pore size.**
Fragment lengths appeared similar across pore sizes (0.2–10.0 µm), though DNA concentration decreased with larger pores. Note that TapeStation reports >60,000 bp as the upper limit, so differences in larger fragments may not be captured.

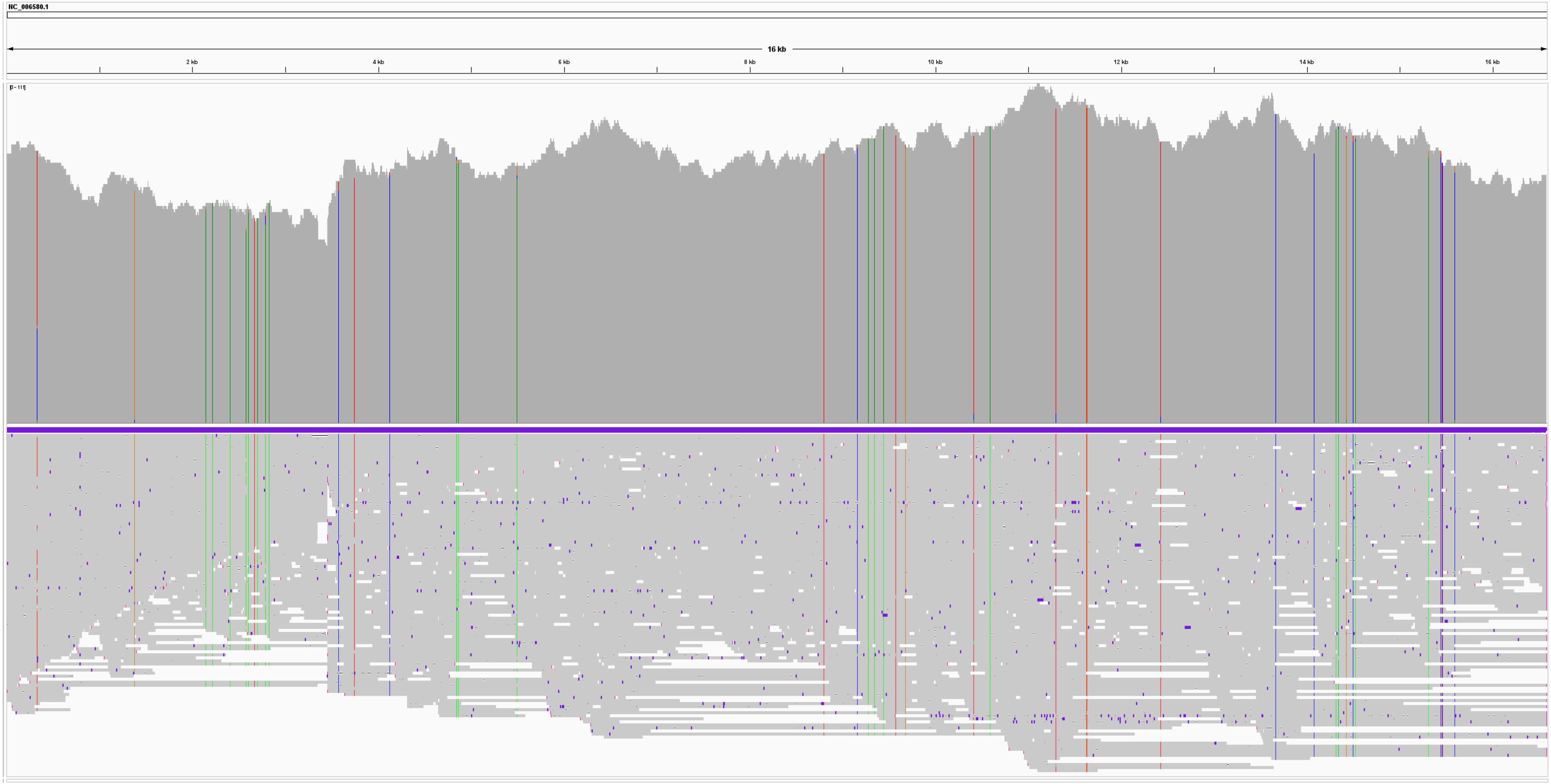


**Supplementary Figure S2. IGV visualization of *C. auratus* reads mapped to mitochondrial reference (sample No.3-6).**
The dark gray horizontal bar at the top indicates the read depth, demonstrating uniform coverage across the entire mitochondrial genome. The occasional vertical lines in red, green, or other colors represent bases recognized as SNVs. Even without PCR amplification, several reads spanned nearly the full mitochondrial genome, showing relatively uniform coverage across its entire length.

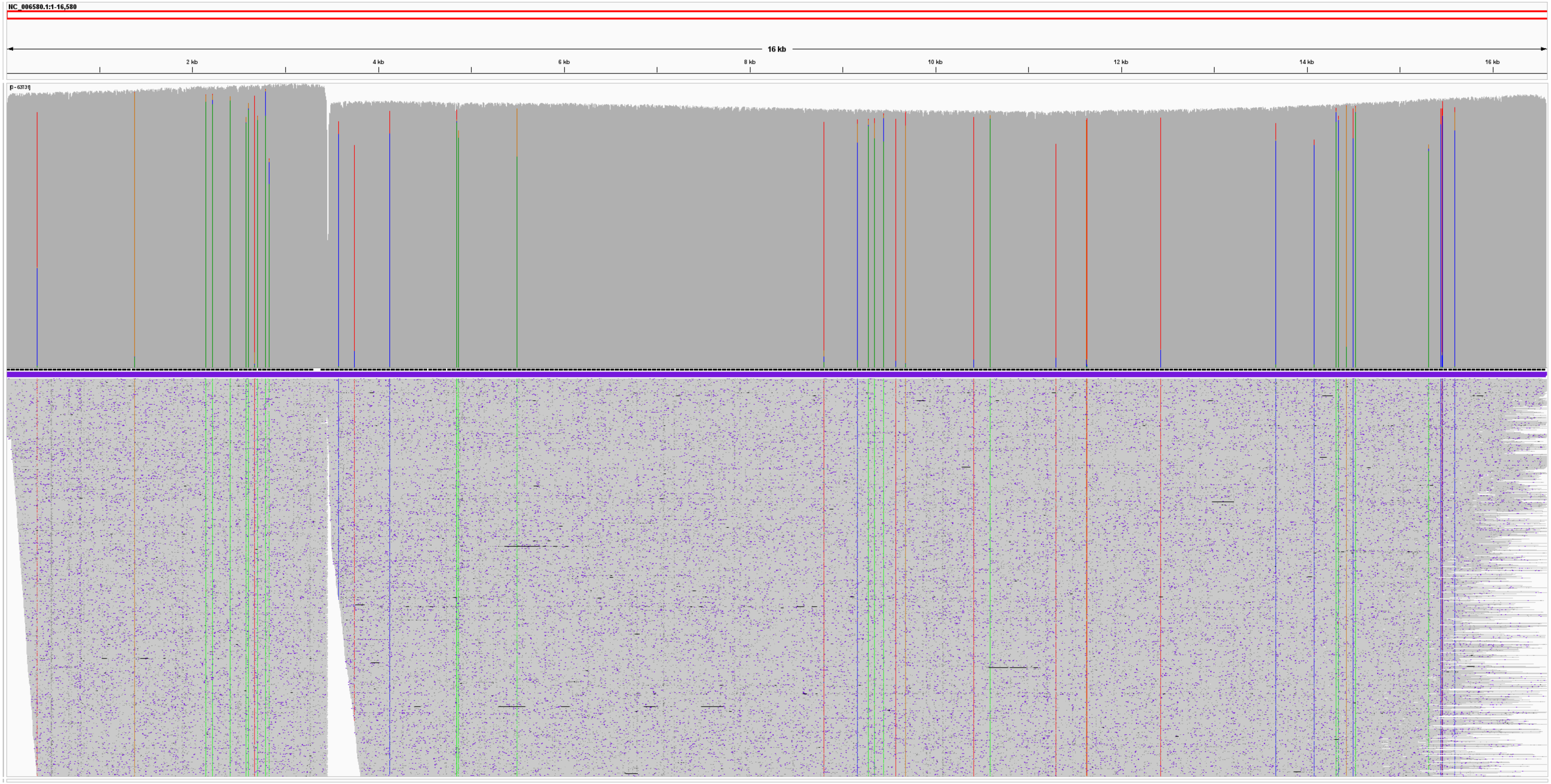


**Supplementary Figure S3. Mapping depth across the mitochondrial genome (sample No.1-2).**
Uniform read depth with high coverage was achieved via the PCR-based approach, although regions with reduced depth correspond to the primer binding sites.

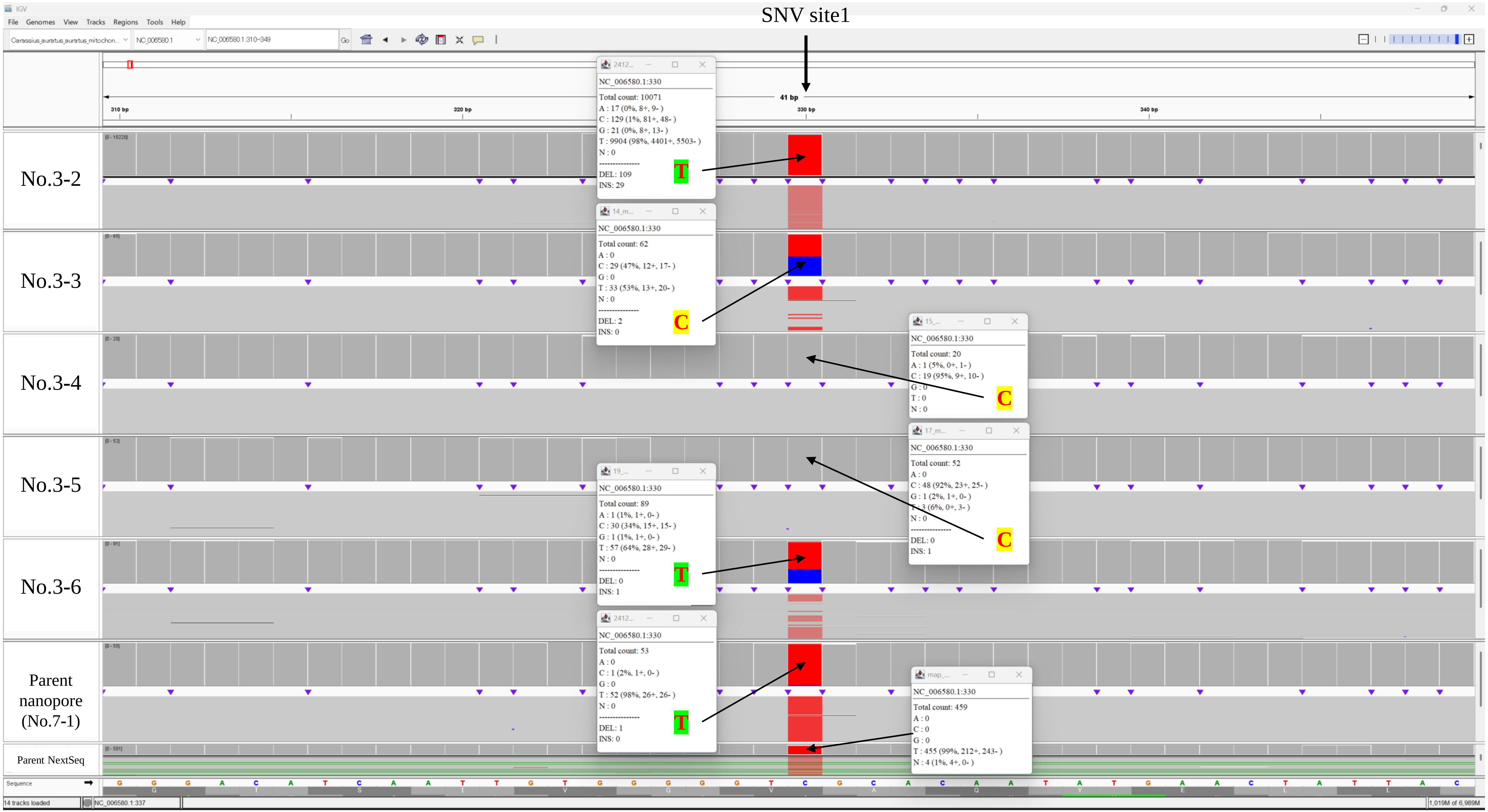


**Supplementary Figure S4. Read alignment around SNV site1.**
In samples No.3-3 and No.3-6, a high proportion of reads carried alleles that differed from those in the assembled sequence.

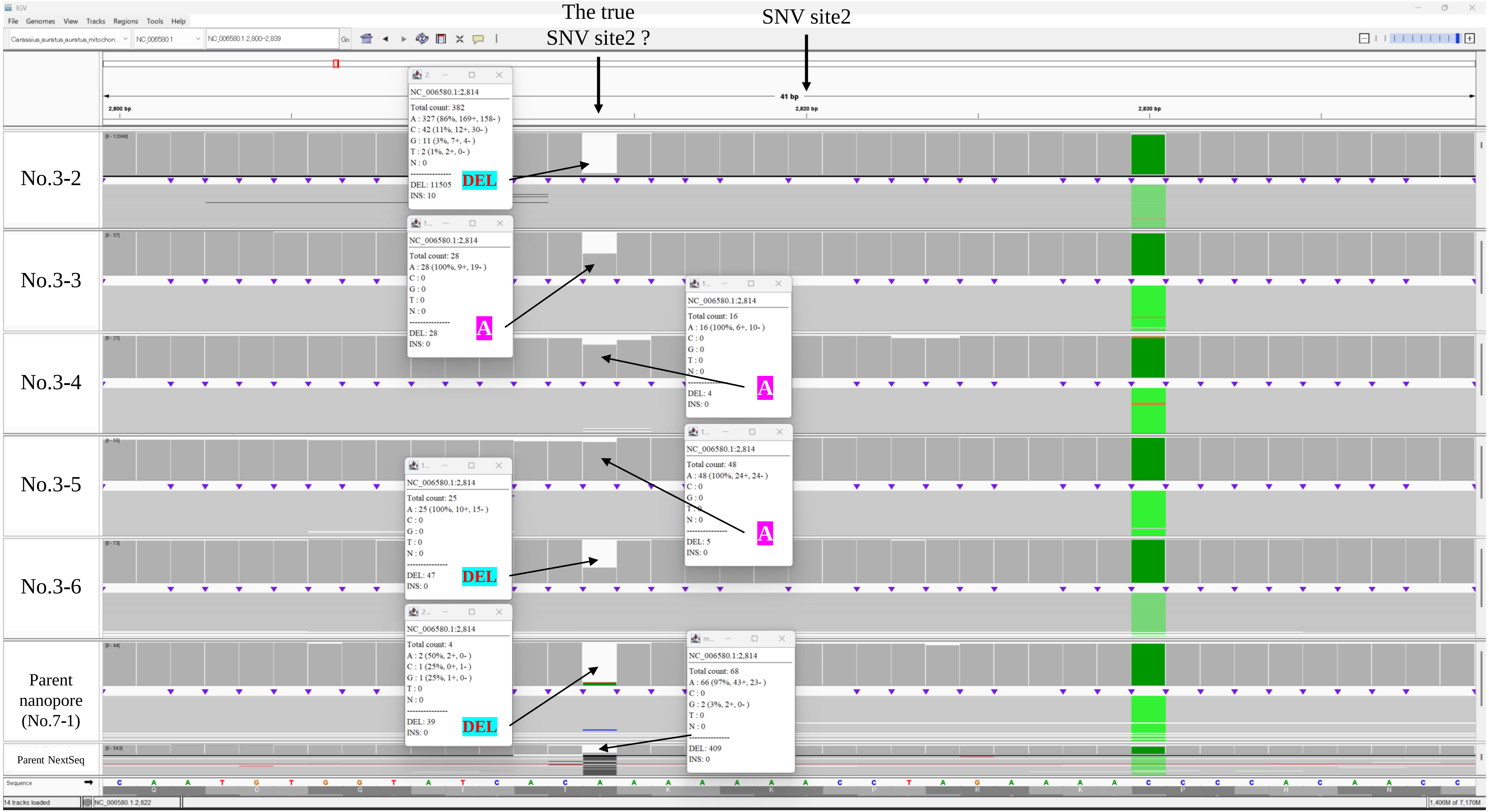


**Supplementary Figure S5. Read alignment around SNV site2.**
In sample No.3-3, the assembled sequence indicated an “A” allele, whereas approximately 50% of the reads exhibited a deletion genotype at the same site.

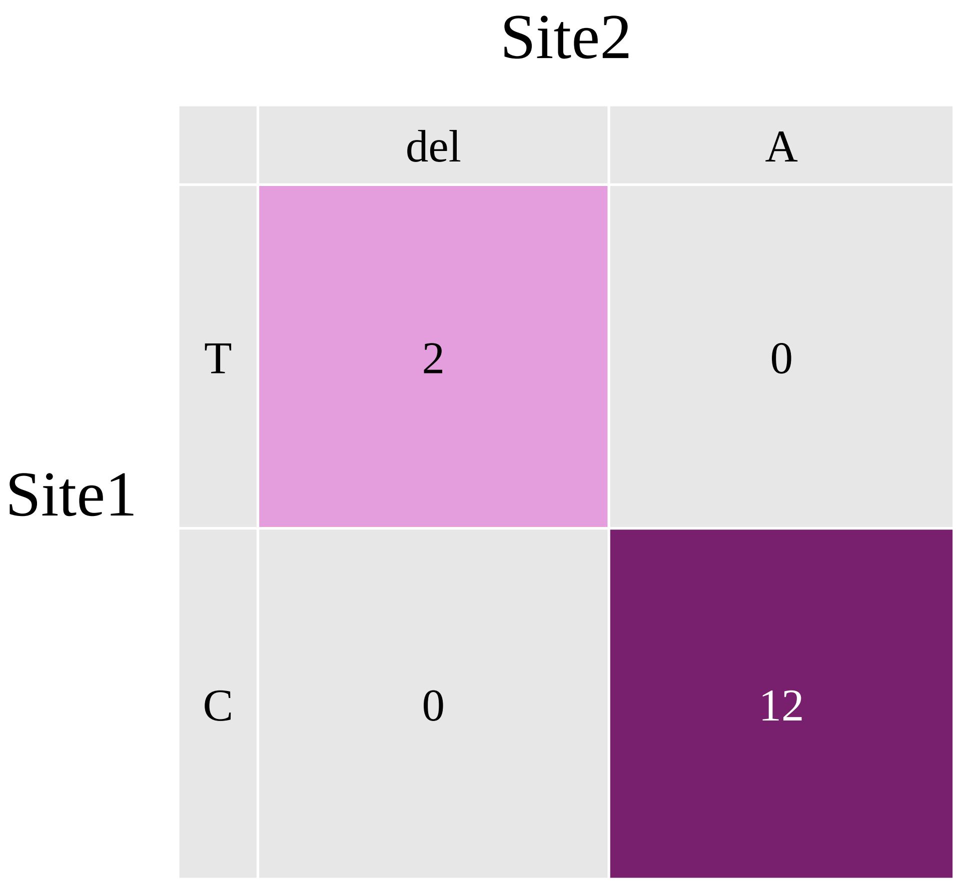


**Supplementary Figure S6. Haplotypes from long reads spanning both SNV site1 and site2 in sample No.3-6.**
Two major haplotypes were reconstructed from single reads exceeding 16,000 bp, indicating the feasibility of detecting linked variants.

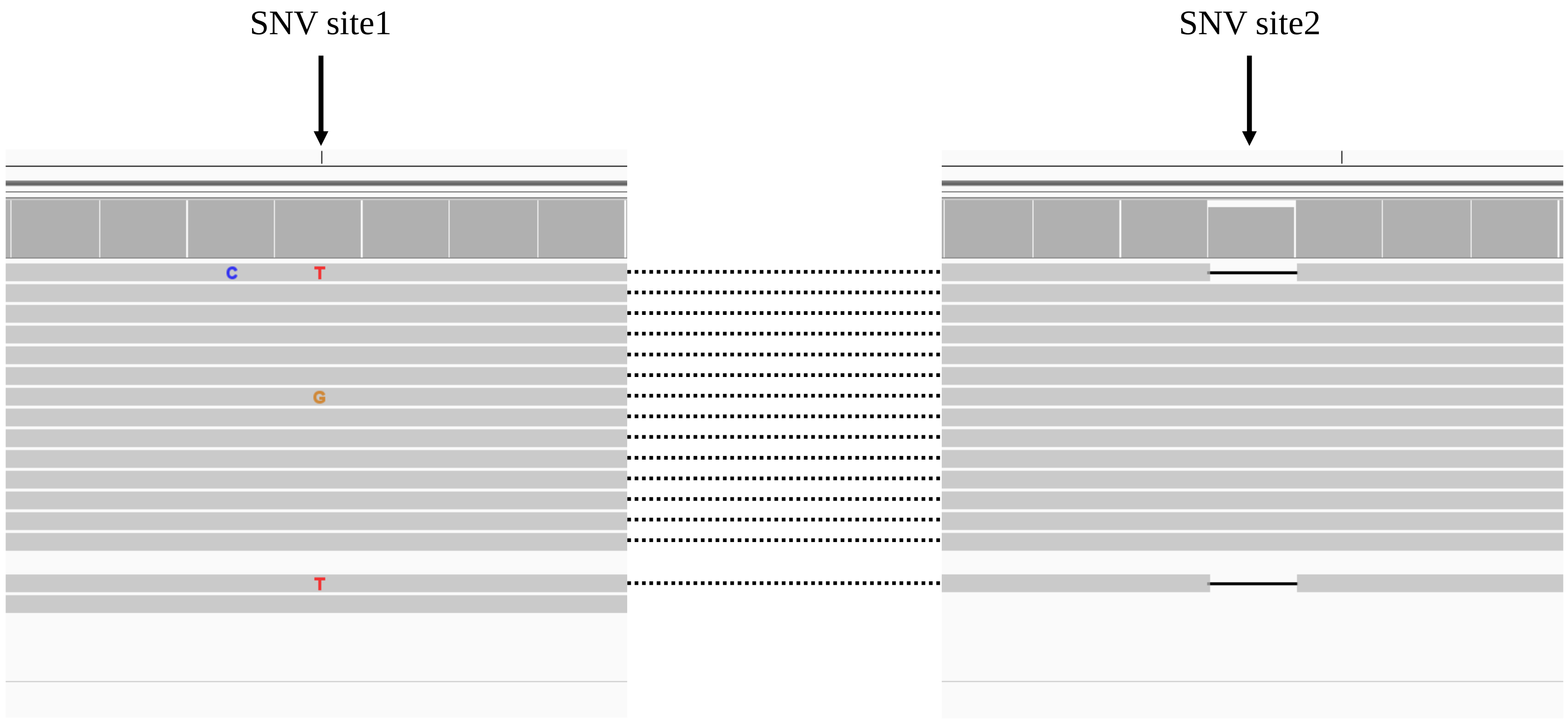


**Supplementary Figure S7. Long-read haplotype inference from sample No.3-6.**
Visualization of reads longer than 16,000 bp that span both Site 1 and Site 2, confirming the mapping patterns and supporting the presence of two major haplotypes.
