## Supplementary Tables for "Noninvasive reconstruction of complete mitochondrial genomes from aquatic environmental DNA using PCR-free long-read sequencing"

**Supplementary Table S1.** **Sampling overview by pore size and sequencing results**

| Spiecies | Pore size | Sequencing Date | Water Sampling Volume (mL) | Total Reads | Mapped Reads | Mapping Rate (%) |
| --- | --- | --- | --- | --- | --- | --- |
| *C. Auratus*  (NCBI RefSeq; GCF_003368295.1) | 0.2 | 2024/6/21 | 250 | 143,479 | 5,398 | 3.8 |
|  | 0.8 | 2024/6/21 | 250 | 255,791 | 20,948 | 8.2 |
|  | 1.2 | 2024/6/21 | 250 | 227,294 | 22,249 | 9.8 |
|  | 2.0 | 2024/6/21 | 250 | 378,156 | 33,690 | 8.9 |
|  | 3.0 | 2024/5/20 | 250 | 213,022 | 18,087 | 8.5 |
|  | 5.0 | 2024/6/21 | 250 | 483,185 | 19,289 | 4.0 |
|  | 10.0 | 2024/6/21 | 250 | 390,271 | 10,334 | 2.6 |
|  | 0.2 | 2024/7/11 | 250 | 64,629 | 3,764 | 5.8 |
|  | 0.8 | 2024/7/11 | 250 | 156,473 | 7,233 | 4.6 |
|  | 1.2 | 2024/7/11 | 250 | 130,445 | 11,433 | 8.8 |
|  | 2.0 | 2024/7/11 | 250 | 124,934 | 12,068 | 9.7 |
|  | 3.0 | 2024/7/11 | 250 | 125,594 | 3,875 | 3.1 |
|  | 5.0 | 2024/7/11 | 250 | 245,086 | 7,434 | 3.0 |
|  | 10.0 | 2024/7/11 | 250 | 110,906 | 4,743 | 4.3 |
|  | 0.2 | 2024/9/6 | 500 | 27,060 | 1,543 | 5.7 |
|  | 0.8 | 2024/9/6 | 500 | 54,588 | 4,605 | 8.4 |
|  | 1.2 | 2024/9/6 | 500 | 78,884 | 9,187 | 11.6 |
|  | 2.0 | 2024/9/6 | 500 | 72,536 | 7,009 | 9.7 |
|  | 3.0 | 2024/9/6 | 500 | 150,388 | 12,448 | 8.3 |
|  | 5.0 | 2024/9/6 | 500 | 113,091 | 8,117 | 7.2 |
|  | 10.0 | 2024/9/6 | 500 | 131,411 | 5,876 | 4.5 |
| *O. Latipes*  (NCBI RefSeq; GCF_002234675.1) | 0.2 | 2024/9/27 | 500 | 112,437 | 1,326 | 1.2 |
|  | 0.8 | 2024/9/27 | 500 | 117,728 | 1,597 | 1.4 |
|  | 1.2 | 2024/9/27 | 500 | 75,825 | 1,474 | 1.9 |
|  | 2.0 | 2024/9/27 | 500 | 135,810 | 1,898 | 1.4 |
|  | 3.0 | 2024/9/27 | 500 | 177,528 | 1,686 | 0.9 |
|  | 5.0 | 2024/9/27 | 500 | 167,296 | 1,548 | 0.9 |
|  | 10.0 | 2024/9/27 | 500 | 119,262 | 1,125 | 0.9 |
|  | 0.2 | 2024/9/27 | 500 | 67,599 | 1,151 | 1.7 |
|  | 0.8 | 2024/9/27 | 500 | 95,632 | 2,733 | 2.9 |
|  | 1.2 | 2024/9/27 | 500 | 78,792 | 3,417 | 4.3 |
|  | 2.0 | 2024/9/27 | 500 | 109,899 | 4,634 | 4.2 |
|  | 3.0 | 2024/9/27 | 500 | 163,259 | 3,445 | 2.1 |
|  | 5.0 | 2024/9/27 | 500 | 127,826 | 2,322 | 1.8 |
|  | 10.0 | 2024/9/27 | 500 | 124,482 | 2,300 | 1.8 |
|  | 0.2 | 2024/9/27 | 500 | 52,211 | 952 | 1.8 |
|  | 0.8 | 2024/9/27 | 500 | 86,410 | 2,792 | 3.2 |
|  | 1.2 | 2024/9/27 | 500 | 76,662 | 2,925 | 3.8 |
|  | 2.0 | 2024/9/27 | 500 | 132,972 | 2,914 | 2.2 |
|  | 3.0 | 2024/9/27 | 500 | 153,784 | 2,293 | 1.5 |
|  | 5.0 | 2024/9/27 | 500 | 198,548 | 1,577 | 0.8 |
|  | 10.0 | 2024/9/27 | 500 | 202,099 | 819 | 0.4 |

**Supplementary Table S2. Two-locus polymorphisms in each assembly-derived sequence of *C. auratus***

|  | Site1 (pos;330, D-loop) | Site2 (pos;2820, 16S rRNA) |
| --- | --- | --- |
| 3-2 | T | - |
| 3-3 | C | A |
| 3-4 | C | A |
| 3-5 | C | A |
| 3-6 | T | - |
| parent | T | - |
| reference | C | A |
